# Exponent Polytopes for Abstracting Chemical Reaction Networks into Formal Specification Models for Compositional Design of Biomolecular Circuits

**DOI:** 10.64898/2026.11.13.766398

**Authors:** Ayush Pandey

## Abstract

Model based design has shown promise for synthetic biological circuits. However, many approaches remain empirical and limited in scale. Here, we develop specification models from chemical reaction network (CRN) models. These specifications encode assumptions explicitly and guarantees that must be met under those assumptions. Specifically, we show how monomial dominance relations in open CRNs induce polyhedral regime assumptions in logarithmic coordinates. These assumptions give polyhedral assume guarantee contracts that encode system behavior. Further, we show how order of magnitude dominance assumptions can be written explicitly as contract assumptions and used to derive tighter guarantees. We demonstrate two biological circuit design applications: identifying the biological mechanisms that meet the required system-level specification, and screening promoter and ribosome binding site choices to determine expression gains that satisfy a specification through contract composition and refinement. Ultimately, these results provide a path from CRN descriptions of synthetic biological parts to specification-based compositional design.

## I. Introduction

Many synthetic biology advances have been driven by mathematical modeling, design automation, and control theory tools [2], [3], [4]. Heuristic design where scientists make decisions based on experience together with model based analysis is common. In practice, circuit designs are often studied by building ordinary differential equation (ODE) models and running simulations to test hypotheses. Models can support useful predictions about fixed points, oscillations, resource competition, and inducer levels needed for desired behavior [5]. However, this does not scale well with the number of components. On the other hand, from a design automation perspective [6], significant progress has been made in designing larger systems that achieve desired logical behavior. For example, Cello [7] compiles a Verilog based specification into biocircuit implementations using libraries of characterized logic gates. Despite this progress, biological design pipelines still cannot readily account for context dependence to achieve desired dynamic properties.

To achieve desired properties and output levels in a synthetic biological circuit, designers often explore various combinations of biological components and their parameters. For example, to express a gene at certain levels, the choices may involve selecting appropriate promoters, ribosome binding sites (RBSs), and regulatory elements. Designers can choose among these circuit parts by running library screening experiments. In some cases, simulations of computational models can be used to reject combinations of parts and parameters that are unlikely to meet the desired specifications [8]. Further along these lines, reinforcement learning and related optimization methods have increasingly been used to automate this process by iteratively testing candidate designs, rejecting poor performing choices, and searching for designs that optimize a specified cost function [8], [9]. When coupled with automated liquid handling systems, these approaches can support largely automated design, build, test, and learn workflows in biofoundries [10]. Despite these advances, a fundamental cost remains: evaluating candidate designs involves time and resource intensive experiments and measurements, or computationally expensive simulation of detailed models of the full circuit.

A related challenge arises when reduced models are used for design. Assumptions such as excess of a species, saturation [11], or time scale separation [12] are implicitly used in deriving such models. As a result, a model may correctly capture a subsystem in one operating regime, but it is difficult to use compositionally because the conditions under which the model is valid are not written explicitly. Consequently, it is difficult to determine whether a composed design of multiple subsystems will work as expected. Monomial dominance and order of magnitude constraints have also been used to derive reduced biochemical models through tropical equilibration [13]. Such conditions do not fully establish whether the operating assumptions of each subsystem in the design remains satisfied under compositon. This motivates the use of formal methods for design.

Formal methods have been used to specify and verify synthetic biological circuits using temporal logic and finite abstraction approaches [14], [15]. Recently, quantifier elimination using assume guarantee contract algebra was used to derive parameter constraints for a class of monotone equilibrium input and output specifications, with applications to biomolecular circuits subject to resource competition and internal feedback [16]. Further, assume guarantee contracts have been used to express circuit design goals and guide refinement from abstract specifications to detailed mathematical models [17]. Although designing systems starting from specifications is the norm in conventional engineering, the field of synthetic biology has not matured to a point where this would be possible. In practice, scientists often design biological circuits using available components with mechanistic models and prior experimental characterization. This motivates the reverse problem: deriving component specifications from these models for compositional design. So, the preliminary work in [17] is not directly applicable to this reverse problem. We study this problem by constructing polyhedral contracts directly from mechanistic chemical reaction network (CRN) descriptions and their operating regime assumptions. These contracts connect the conditions used in reduced models to compositional verification.

We use these contracts for compositional verification against system level specifications, with applications to mechanism selection and biological part screening. The specific contributions of this paper are given below:

### Main contributions

1. Using CRN exponent polytopes [18], we construct polyhedral assume guarantee contracts [19] for a general class of positive quotient CRNs. The assumptions in these contracts explicitly express commonly used modeling assumptions: monomial dominance, order-of-magnitude dominance, reaction balances, and conservation laws. We derive corresponding guarantees to develop these specification models.
2. For commonly used Hill propensities in CRNs, we derive assume-guarantee contracts that include input dependent guarantees. We apply this to construct a library of contract models for biological parts from the Cello [7] logic gate library.
3. We apply our results to two specific biological circuit design tasks: (a) selecting mechanisms that satisfy an AND specification, and (b) screening promoter and RBS combinations from a library to find the combination of parts that achieve the desired specification.

This paper extends our CDC 2026 conference paper [1] where we established the construction of assume-guarantee contracts from monomial dominance and exponent polytopes for mass action CRNs only. Here, we provide the full proofs, add new equilibrium balance and refinement results, together with the library construction and design applications described above. In this paper, we also demonstrate the accuracy of predictions for the biological circuit applications using previously published experimental data.

## II. Problem Formulation

We present the compositional design and analysis problem at the end of this section. To set it up, we first need to define CRN subsystems and their propensity classes, polyhedral contracts, conservation laws, and equilibrium behavior.

### Definition 1

(CRN subsystem). A CRN consists of a finite species set *S*, parameters *θ*, and a finite reaction set 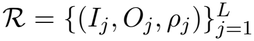, where *I*_*j*_, *O*_*j*_ are multisets of input and output species and *ρ*_*j*_ is a nonnegative propensity function [20]. Among the species, we label the internal concentrations as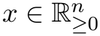, inputs as 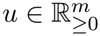, and we denote the outputs by *y* = *h*(*x, u*) . Let *α*_*j*_, *β*_*j*_ be the internal and external reactant stoichiometric vectors and 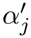 the internal product stoichiometric vector. We define the net stoichiometric change 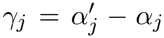 the stoichiometric matrix

*N* = [*γ*_1_· · ·*γ*_*L*_], and the reaction rate *r*_*j*_(*x, u*) = *ρ*_*j*_(*x, u*; *θ*). Then, the subsystem dynamics are 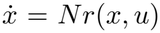. We denote the subsystem by Σ = (*x, u, y*).

Note that we model reversible reactions as two separate irreversible reactions.

### Definition 2

(Positive quotient CRN). We define a positive quotient CRN as one with propensities *r*_*j*_(*x, u*) = *A*_*j*_(*x, u*)*/B*_*j*_(*x, u*), where *A*_*j*_, *B*_*j*_ are finite positive generalized monomial sums, where each monomial has the form *cw*^*a*^ with *c >* 0, *w >* 0, and a fixed real exponent vector *a*.

Next, we present two sub-classes of CRNs.

### Definition 3

(Mass action CRN). We define a mass-action CRN with propensities 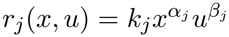, where *k*_*j*_ are positive rate constants. That is, 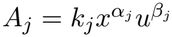 and *B*_*j*_ = 1.

### Definition 4

(Hill propensity CRN). We define a Hill propensity CRN as one that has reactions with either mass-action propensities or Hill propensities of one of the forms in Table 1. The Hill propensities have the general form

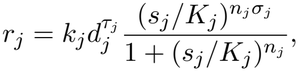

where *k*_*j*_, *K*_*j*_, *n*_*j*_ *>* 0, *s*_*j*_ is the regulating species concentration, and *d*_*j*_ is the species concentration multiplying the proportional rate. For these propensities, 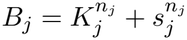.

**TABLE I.** Standard Hill function forms [20].

| Form | $\sigma_j$ | $\tau_j$ | $A_j$ | $r_j = A_j/B_j$ |
|---|---|---|---|---|
| Positive | 1 | 0 | $k_j s_j^{n_j}$ | $\frac{k_j s_j^{n_j}}{K_j^{n_j} + s_j^{n_j}}$ |
| Negative | 0 | 0 | $k_j K_j^{n_j}$ | $\frac{k_j K_j^{n_j}}{K_j^{n_j} + s_j^{n_j}}$ |
| Proportional positive | 1 | 1 | $k_j d_j s_j^{n_j}$ | $\frac{k_j d_j s_j^{n_j}}{K_j^{n_j} + s_j^{n_j}}$ |
| Proportional negative | 0 | 1 | $k_j d_j K_j^{n_j}$ | $\frac{k_j d_j K_j^{n_j}}{K_j^{n_j} + s_j^{n_j}}$ |

We write the dynamics in production and degradation form by collecting the positive and negative stoichiometric contributions to each state coordinate [21],

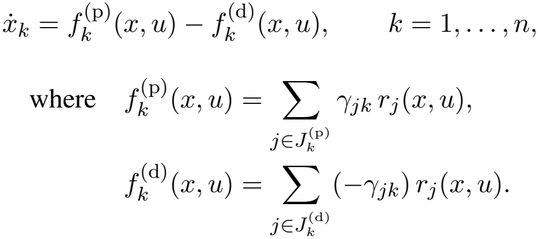

for 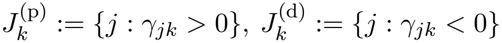. For mass action kinetics, we introduce the notation 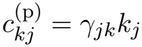 and 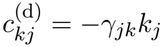 When the context is clear, we drop the superscripts and write *c*_*kj*_ for the positive coefficients. Next, at a higher level abstraction, we define specification models for a subsystem.

### Definition 5

(Polyhedral assume guarantee contract). For a subsystem Σ, let *ξ* denote selected abstraction variables, which may include states, inputs, outputs, or auxiliary variables. A polyhedral assume guarantee contract is a pair *C* = (*a, g*) with

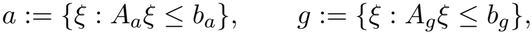

where *a* specifies the admissible operating conditions and *g* specifies the properties required under those conditions. We write Σ |= C if, pointwise, whenever the subsystem variables satisfy *a*, they also satisfy *g*.

### A. Conservation Laws and Equilibrium Behavior

Following standard CRN literature, we represent conservation laws by a matrix 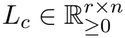, which has independent rows and satisfies *L N* = 0. The conserved total resources are 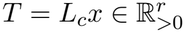 These totals remain constant along the subsystem dynamics because 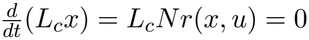. With *v* = (*u, T* ), we define the positive equilibrium set *ℰ* (*v*) = {*x >* 0 : *Nr*(*x, u*) = 0, *L*_*c*_*x* = *T* }.

#### Assumption 1.

We assume that the CRN has at least one positive equilibrium, that is, *ℰ* (*v*) ≠ *∅* for every *v ∈V*, where *V* is the given set of admissible values of the inputs *u* and conserved total resources *T* .

We state a sufficient condition for Assumption 1 next.

#### Proposition 1.

*Suppose f* (*x, u*) = *Nr*(*x, u*) *is continuous on the set of positive states satisfying the conservation laws L*_*c*_*x* = *T* . *For each v* = (*u, T* ) *∈V, let there be finite concentration bounds* 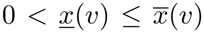 *such that at least one state satisfies the conservation laws and lies within these concentration bounds. That is*, 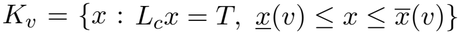 *is nonempty. If each component of f* (*x, u*) *is nonnegative at its corresponding lower concentration bound and nonpositive at its upper concentration bound on K*_*v*_, *then Assumption 1 holds. The same conclusion holds if all these boundary signs are reversed*.

*Proof*. Fix *v* = (*u, T* ) *∈ V* and consider the continuous map from the states *x ∈ K*_*v*_ to the point of *K*_*v*_ nearest to *x* + *f* (*x, u*). Since *K*_*v*_ is nonempty, compact, and convex, the fixed point theorem [22] gives a fixed point *x*^*\**^, so *L*_*c*_*f* (*x*^*\**^, *u*) = 0. Moreover, *f*_*i*_(*x*^*\**^, *u*) ≥ 0 when 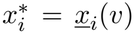 and *f*_*i*_(*x*^*\**^, *u*) ≤ 0 when 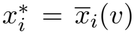. Thus *x*^*\**^ + *sf* (*x*^*\**^, *u*) *∈ K*_*v*_ for all sufficiently small *s >* 0. If *f* (*x*^*\**^, *u*) ≠ 0, choosing such an *s* 1 gives a point in *K*_*v*_ whose distance to *x*^*\**^ + *f* (*x*^*\**^, *u*) is (1−*s*) *∥f* (*x*^*\**^, *u*) *∥ < ∥f* (*x*^*\**^, *u*) *∥* . This contradicts the choice of *x*^*\**^ as the nearest point. Therefore *f* (*x*^*\**^, *u*) = 0, and the positive lower concentration bounds give *x*^*\**^ *>* 0. Since *v* was arbitrary, Assumption 1 holds. If the boundary signs are reversed, the same argument can be applied to −*f* .

We note that the above holds for a broad class of biomolecular systems. Some examples include Hill regulation models with bounded, strictly positive production and linear degradation, simple reversible binding systems with positive conserved total resources, and systems modeled as birth-death processes. In addition, if uniqueness is desired, we can impose injectivity of the vector field on each conservation class.

### B. Contracts for Equilibrium Behavior

For positive outputs *y* = *h*(*x, u*), we define logarithmic coordinates *z*_*x*_ := log *x, z*_*v*_ := log *v*, and *z*_*y*_ := log *y*. For equilibrium behavior, we specialize Definition 5 to C = (*a, g*),

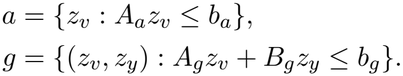

We write Σ |= *C* if, for every *z*_*v*_ *∈ a*, E(*v*) is nonempty and every *x ∈ ℰ* (*v*) satisfies (*z*_*v*_, log *h*(*x, u*)) *∈ g*.

For contracts with a common interface, we use the refinement relation of [23]. A contract *C*_1_ = (*a*_1_, *g*_1_) refines *C*_2_ = (*a*_2_, *g*_2_), written *C*_1_ *⪯ C*_2_, when

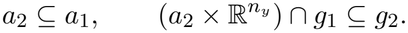

### C. Problem: Compositional Design and Analysis

The key problem we study in this paper is compositional design and analysis of biological circuits. That is, given open CRN subsystems Σ_1_, …, Σ_*N*_ and a top-level system specification *C*_spec_, our goal is to derive polyhedral contracts *C*_1_, …, *C*_*N*_ that describe subsystem behavior under explicit operating assumptions. Then, using composition and refinement of these contracts, we aim to determine whether Σ_full_ = Σ_1_ *∥* · · · *∥* Σ_*N*_ satisfies *C*_spec_ and to identify component and mechanism choices that meet the specification.

## III. Preliminaries

For mass action CRNs, each production or degradation term is a positive sum of monomials in the variables (*x, u*). We write this family of monomials with positive coefficients over a finite index set *I*, as

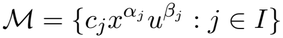

and the sum of these monomials as

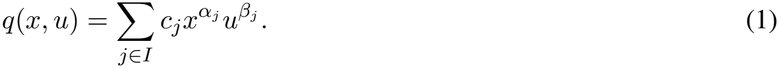

For each monomial in ℳ, we define the exponent set as *E*(ℳ) := { (*α*_*j*_, *β*_*j*_) : *j ∈ I*}, and the associated exponent polytope is

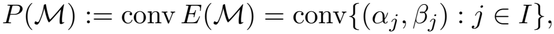

where conv denotes the convex hull. We have positive generalized monomials 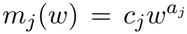, where *w >* 0, *c*_*j*_ *>* 0, and *a*_*j*_ are fixed real exponent vectors.

To distinguish between different monomials in the family, we introduce the notion of dominance.

### Definition 6

(Monomial dominance and region). For a monomial family ℳ, we say that *m*_*j*_ dominates *m*_*ℓ*_ at *w* if *m*_*j*_(*w*) ≥ *m*_*ℓ*_(*w*). More generally, *m*_*j*_ dominates *m*_*ℓ*_ with margin *η >* 0 if *m*_*j*_(*w*) ≥ *ηm*_*ℓ*_(*w*). A region is a nonempty set given by finitely many such relations. For mass action CRNs, *w* = (*x, u*) and 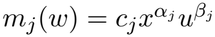.

With this notion of dominance, we can now establish the first lemma which justifies the use of logarithmic coordinates in the analysis of the CRN dynamics. For the logarithmic analysis, we restrict attention to the strictly positive orthant. The following characterization follows from [18].

### Lemma 1.

*For* 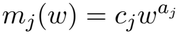 *and z* = log *w, the relation m*_*j*_(*w*) ≥ *ηm*_*ℓ*_(*w*) *is equivalent to*

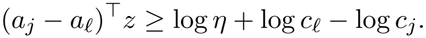

*Consequently, every region is polyhedral in logarithmic coordinates*.

*Proof*. Taking logarithms of the two positive monomials and subtracting gives the stated inequality.

To derive contract specifications from the CRN model, we start by defining various operating regimes.

### Definition 7

(Regime). For the subsystem Σ, a regime is a nonempty subset of 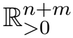 specified by a finite collection of pairwise dominance relations among the monomials appearing in *f* ^(p)^(*x, u*) and *f* ^(d)^(*x, u*).

The definition immediately implies the following result.

### Lemma 2.

*Any regime defined by finitely many monomial dominance relations is polyhedral in* (*z*_*x*_, *z*_*u*_).

To connect the positive sums of monomials in equation (1) to convex polytopes required by polyhedral contracts (Def. 5), we use the following result from [18].

### Theorem 1.

*For the positive monomial sum*

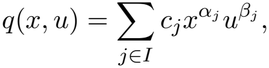

*its logarithmic derivative vector satisfies*

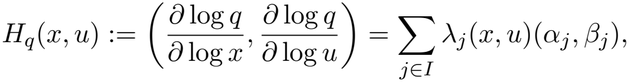

*where*

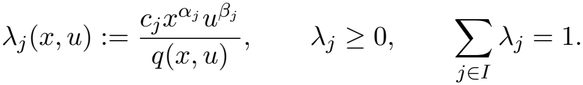

*Hence, H*_*q*_(*x, u*) *∈ P* (ℳ). *The identity also holds for* 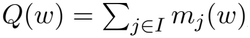 *with w and a*_*j*_ *in place of* (*x, u*) *and* (*α*_*j*_, *β*_*j*_).

*Proof*. For each state coordinate, differentiating the positive monomial sum gives

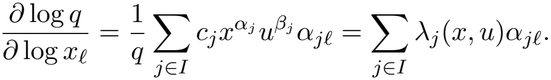

The same calculation for each input coordinate gives *∂* log *q/∂* 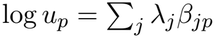. Stacking these coordinates gives the stated vector expression for *H*_*q*_. Since the coefficients are nonnegative and sum to one, *H*_*q*_ lies in the convex hull of the exponent vectors.

Applying Theorem 1, for *s ∈* {p, d} and 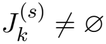,

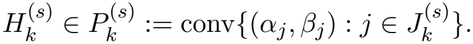

## IV. Polyhedral Contracts for Chemical Reaction Networks

### Theorem 2.

*For every regime ℛ of a deterministic mass action CRN, there exists a polyhedral assume guarantee contract C*_*ℛ*_ = (*a*_*ℛ*_, *g*_*ℛ*_), *where*

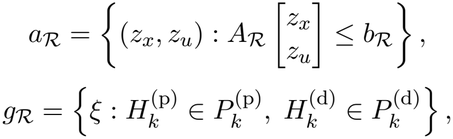

*with each constraint imposed whenever the corresponding reaction set is nonempty. Moreover*, Σ |= *C*_*ℛ*_.

*Proof*. By Lemma 2, *a*_*R*_ is polyhedral. Theorem 1 places each logarithmic derivative in its corresponding exponent polytope whenever the reaction set is nonempty. Thus *g*_*ℛ*_ is polyhedral and holds throughout *a*_*R*_, so Σ |= *C*_*ℛ*_.

### A. Polyhedral contracts under dominance margins

Order of magnitude assumptions such as excess of a species or saturation are commonly used to obtain reduced models [24]. However, these assumptions are often used implicitly. We capture these explicitly in specifications by evaluating monomial dominance margins in CRNs. The results below are for the production terms, but an analogous result holds for the degradation terms. As such, we drop the (p) superscript from all stoichiometric coefficients while keeping it on the reaction sets.

#### Theorem 3.

*Suppose that for a regime ℛ of a deterministic mass action CRN there is a nonempty dominant subset* 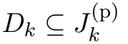 *for all k, such that*

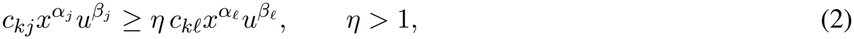

*for all j∈D*_*k*_ *and* 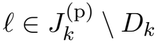. *Then there exists a polyhedral assume guarantee* 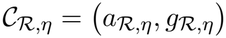 *where the assumption set is obtained by conjoining the logarithmic form of the dominance inequalities to the regime inequalities*,

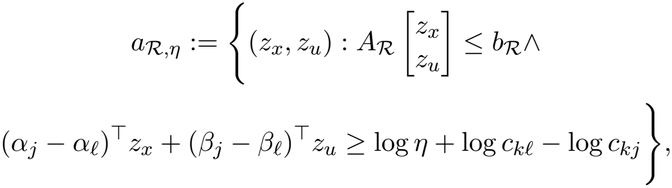

*and the guarantee set is the baseline logarithmic derivative inclusion conjoined with the additional nondominant weight bound*

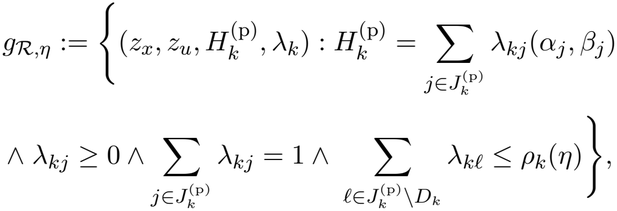

*with constants*

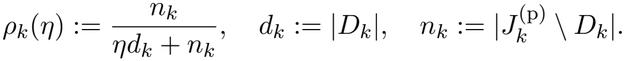

*Moreover* 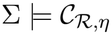.

*Proof*. By Lemma 1, the dominance condition (2) is equivalent to the affine inequality in the assumption set. From Theorem 1, we have the baseline convex combination relation for the logarithmic derivative vector, here, 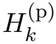. It remains to bound the total weight assigned to the nondominant monomials. Define the dominant and nondominant sums as

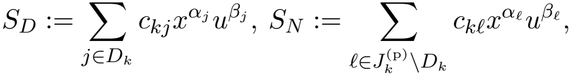

then, the nondominant weight is

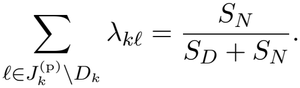

We rewrite the dominance assumption as 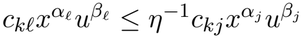 and sum this inequality over all dominant and nondominant indices to get

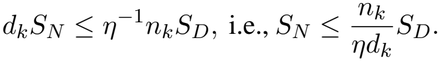

Substituting into the weight expression yields

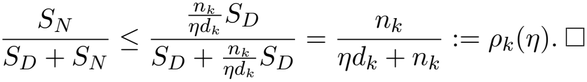

### B. Monotonicity from sensitivity polytopes

The preceding results give polyhedral guarantees on logarithmic sensitivities. We use these guarantees to establish whether a positive reaction rate or interface function increases or decreases with each variable. Once these signs are established throughout a region given by interval bounds, the extrema of the function can be determined by endpoint evaluations. So, consider a positive quotient *F* (*v*) = *P* (*v*)*/Q*(*v*), the identity *H*_*F*_ = *H*_*P*_ − *H*_*Q*_ gives a sensitivity polytope

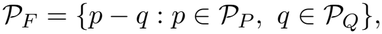

where *P*_*P*_ and *P*_*Q*_ bound the logarithmic sensitivities of the positive sums *P* and *Q*. The following result establishes monotonicity by linear optimization over this polytope.

#### Lemma 3.

*Let F* (*v*) *>* 0 *be continuously differentiable on* 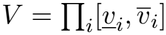 *with positive interval bounds. Suppose a polytope* P_*F*_ *contains the logarithmic sensitivity* 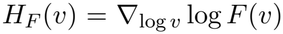 *for every v ∈ V* . *If* 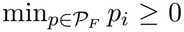 *then F is nondecreasing in v*_*i*_. *If* 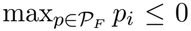, *then F is nonincreasing in v*_*i*_. *When one of these conditions holds for every coordinate, the minimum and maximum of F over V are attained at the corners chosen according to these signs*.

*Proof*. Since

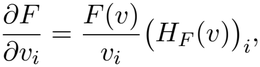

the polytope bounds establish the derivative signs. We can vary each coordinate within the corresponding interval to establish monotonicity. The minimum uses the lower endpoint for a nondecreasing coordinate and the upper endpoint for a nonincreasing coordinate. Similarly, the maximum uses the opposite endpoints of the region.

### C. Numerical example

To interpret these results, consider a production term with three monomials 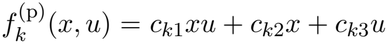, and an order of magnitude dominance relation (*D*_*k*_ = {1, 2}) given by 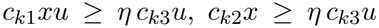, for *η >* 1. Then, from Theorem 1, the baseline guarantee is that the logarithmic derivative vector of 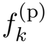 with respect to (*x, u*), denoted by 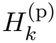 is a convex combination of the exponent vectors of the three monomials: 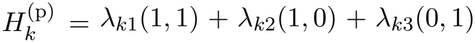 where each *λ*_*kj*_ ≥ 0 and together they sum to one. Theorem 3 gives us an additional guarantee that the total weight assigned to the nondominant monomial, *λ*_*k*3_, is upper bounded by

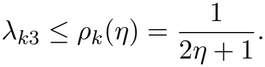

Hence, the contract guarantees tighten to 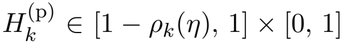. In this example, the dominant exponent set conv{(1, 1), (1, 0)} is one dimensional, that is, a line segment with first coordinate equal to 1. The dominance assumption bounds the contribution of the nondominant monomial (0, 1), so the first coordinate of 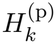 differs from that of the dominant exponent segment by at most *ρ*_*k*_(*η*).

## V. Equilibrium Interface Contracts

Using Theorem 3, we can abstract system assumptions and guarantees using logarithmic sensitivities of CRN production and degradation rates. However, it is also common for circuit specifications to include bounds on equilibrium concentrations of species and output fluxes. So, in what follows, we impose such constraints to derive the guarantees from equilibrium balances and conservation laws for CRNs.

### A. Bounds from monomial ratios

To bound the value of a positive monomial sum, we bound each term relative to a reference monomial and sum the resulting inequalities.

#### Lemma 4.

*For a positive monomial sum* 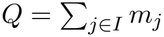 *choose a reference monomial j*_0_ *∈ I and define the lower and upper bounds for each monomial ratio as*

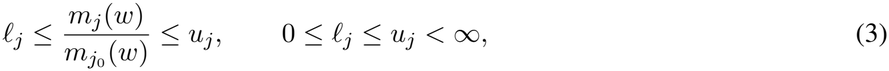

*with* 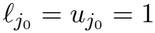. *Then, we have that*

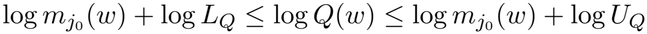

*where* 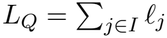, *and* 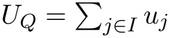.

*Proof*. Divide *Q* by 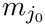, sum (3), and take logarithms. The reference term ensures *L*_*Q*_≥1. Note that both bounds are affine in log(*w*).

### B. Contracts from equilibrium balances

We can use this result to now write contracts for CRNs with equilibrium balances. For a positive quotient CRN, the equilibrium balance for species *i* is

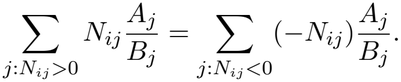

Multiplying this equation by 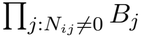 gives a positive generalized monomial sum on each nonzero side. Note that when either side is zero, we impose no equilibrium balance constraint. At equilibrium points *x ∈ ℰ* (*v*) and for positive outputs *y* = *h*(*x, u*), we write the resulting equilibrium balance as *P* (*w*) = *Q*(*w*), where *w* = (*x, u, T, y*). Here *P* and *Q* are finite positive generalized monomial sums.

#### Theorem 4.

*For any positive quotient CRN that satisfies Assumption 1 with equilibrium balances P* (*w*) = *Q*(*w*) *at every equilibrium point, let a nonempty polyhedral assumption, a, specify the set V* = {*v >* 0 : log *v ∈ a*}. *If assumptions establish a region ℛ of finite monomial ratio bounds for the sums P, Q such that w ∈ ℛ for every v ∈ ℰV and x* (*v*), *then the contract C* = (*a, g*) *certifies the output guarantee at every positive equilibrium, where*

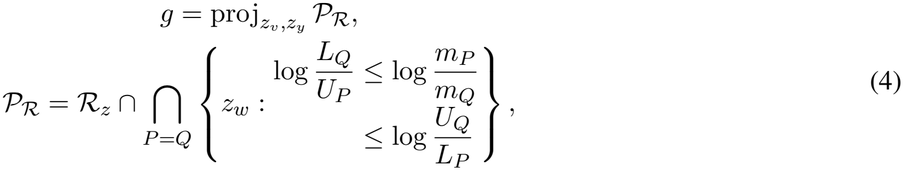

*where L*_*P*_, *U*_*P*_, *L*_*Q*_, *U*_*Q*_ *are constants, ℛ*_*z*_ = {log *w* : *w ∈*ℛ}, *and m*_*P*_, *m*_*Q*_ *are the reference monomials for each balance*.

*Proof*. Fix *v ∈ V* and choose any *x*^*\**^ *∈ ℰ* (*v*), using Assumption 1. Set *y*^*\**^ = *h*(*x*^*\**^, *u*) and *w*^*\**^ = (*x*^*\**^, *u, T, y*^*\**^). Since *w*^*\**^ *∈* ℛ, the chosen monomial ratio bounds hold at *w*^*\**^. From Lemma 4, we can obtain constants *L*_*P*_, *U*_*P*_ by summing the lower and upper bounds on the ratios of the terms of *P* to *m*_*P*_, giving *L*_*P*_ *m*_*P*_ ≤ *P* ≤ *U*_*P*_ *m*_*P*_ . Similarly, we apply the construction to *Q* to get *L*_*Q*_*m*_*Q*_ ≤ *Q* ≤ *U*_*Q*_*m*_*Q*_. At *w*^*\**^, the balance *P* = *Q* therefore implies *L*_*Q*_*/U*_*P*_ ≤ *m*_*P*_ (*w*^*\**^)*/m*_*Q*_(*w*^*\**^) ≤ *U*_*Q*_*/L*_*P*_ . Taking logarithms and combining these bounds for all equilibrium balances gives log *w*^*\**^ *∈ P*_*R*_ as defined in (4). The set P_*R*_ is a polyhedron, and its projection *g* contains (log *v*, log *h*(*x*^*\**^, *u*)). Since *v* and *x*^*\**^ were arbitrary, *g* holds at every positive equilibrium under *a*. Thus Σ |= *C*.

The above result allows us to write contracts that explicitly account for assumptions imposed by equilibrium balances and conservation laws, and the corresponding guarantees under these assumptions.

### C. Refinement of equilibrium guarantees

We can strengthen the guarantees of the contract *C* = (*a, g*) from Theorem 4 by adding valid equilibrium inequalities to P . Note that for contracts with the same assumptions and interface variables, if we have 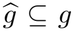 then the new contract, 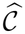, refines the original contract, 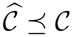 [23]. This motivates the following.

#### Proposition 2.

*Under the conditions of Theorem 4, consider a CRN whose assumptions, a, give additional bounds on the equilibrium balances that are affine in logarithmic coordinates, z*_*w*_ = log *w, and hold at every positive equilibrium under a. Then, the CRN satisfies a contract* 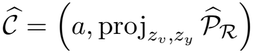 *where* 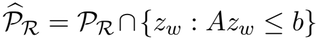 *and Az*_*w*_ ≤ *b collects the additional bounds. The contract satisfies* 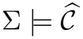 *and* 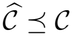.

*Proof*. We can obtain additional affine bounds from the positive balance sums of a CRN. For such a sum 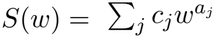, we define *w* = *e*^*z*^ and *F*_*S*_(*z*) = log(*S*(*e*^*z*^)). From Theorem 1, *∂F*_*S*_*/∂z*_*i*_ = *ā*_*i*_, where *ā* = Σ_*j*_ *λ*_*j*_*a*_*j*_ uses the monomial weights of that theorem. Differentiating once more gives

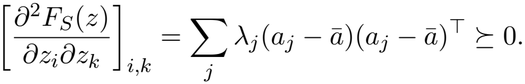

Thus *F*_*S*_ is convex [25], and its tangent at any *z*_0_ gives the affine lower bound

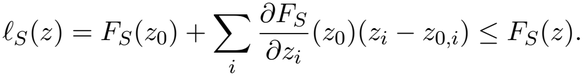

If the additional bounds give a finite polytope *D* for the relevant logarithmic arguments, an affine function *u*_*S*_ with *u*_*S*_(*v*) ≥ *F*_*S*_(*v*) at every vertex *v* of *D* satisfies *F*_*S*_(*z*) ≤ *u*_*S*_(*z*) throughout *D*. This follows by writing *z* as a convex combination of the vertices and using convexity of *F*_*S*_ and affinity of *u*_*S*_. Therefore, a reference monomial can be factored out before this construction, so that *D* bounds only the logarithmic monomial ratios.

For each balance *P* = *Q*, the resulting inequalities *ℓ*_*P*_ ≤ log *P* ≤ *u*_*P*_ and *ℓ*_*Q*_ ≤ log *Q* ≤ *u*_*Q*_ imply the affine equilibrium constraints *ℓ*_*P*_ ≤ *u*_*Q*_ and *ℓ*_*Q*_ ≤ *u*_*P*_ . These constraints can be included in *Az*_*w*_ ≤ *b*. The affine bounds from Lemma 4 can also be combined with these tangent and vertex bounds. In particular, for a positive quotient output *y* = *A*(*u*)*/B*(*u*), they give *ℓ*_*A*_ − *u*_*B*_ ≤ log *y* ≤ *u*_*A*_ − *ℓ*_*B*_. By Theorem 4, every positive equilibrium under *a* has logarithmic coordinates in *P*_*ℛ*_ . It also satisfies the additional bounds, so it belongs to 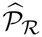. Its interface values therefore belong to 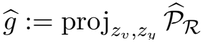, which establishes 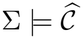. Moreover, 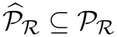 gives *ĝ ⊆ g* under projection, which implies 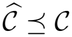.

We note that the guarantees provided by Lemma 4 can be unbounded in logarithmic coordinates, therefore, taking the intersection of those bounds with the inequalities given by Proposition 2 can refine the guarantee under the same assumptions.

### D. Refined contracts for Hill propensities

For Hill propensities, we use Proposition 2 to obtain a refined contract 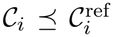, where 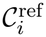 is the contract obtained using Theorem 4 with the monomial ratio bounds on the Hill rate from Lemma 4.

#### Theorem 5.

*For a CRN under the hypotheses of Theorem 4 with Hill propensity r with regulatory species s and d, if input assumptions, a*_*i*_, *give the bounds s ∈* [*s*_*i*_, *s*_*i*+1_] *with* 0 < *s*_*i*_ *<s*_*i*+1_ < ∞ *and positive parameter intervals k ∈* [*k*^*−*^, *k*^+^] *and K ∈* [*K*^*−*^, *K*^+^], *then the CRN satisfies the contract C*_*i*_ = (*a*_*i*_, *g*_*i*_), *which refines* 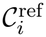. *The contract guarantee g*_*i*_ *can be constructed using*

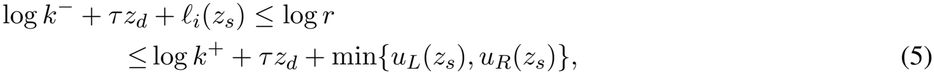

*where z denotes logarithmic coordinates and ℓ*_*i*_ *is the chord over* [log *s*_*i*_, log *s*_*i*+1_] *of the pointwise minimum logarithmic Hill factor over the interval for K, and u*_*L*_, *u*_*R*_ *are the left and right endpoint tangents of the pointwise maximum*.

*Proof*. We can rewrite the Hill forms in Definition 4 as

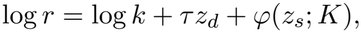

where *φ*(*z*; *K*) = *σn*(*z* − log *K*) − log 1 + *e*^*n*(*z−*log *K*)^ is the only part that is not affine. But we can derive affine bounds on this term using its concavity. With *q* = *e*^*n*(*z−*log *K*)^, direct differentiation gives us

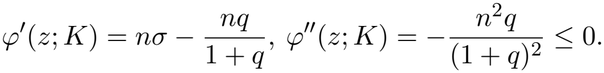

We define *φ*_*−*_(*z*) = min_*K*_ *φ*(*z*; *K*) and *φ*_+_(*z*) = max_*K*_ *φ*(*z*; *K*) over the given interval (activation extrema occur at *K*^+^ and *K*^*−*^, while for repression they occur at *K*^*−*^ and *K*^+^). Then, the chord and endpoint tangents are

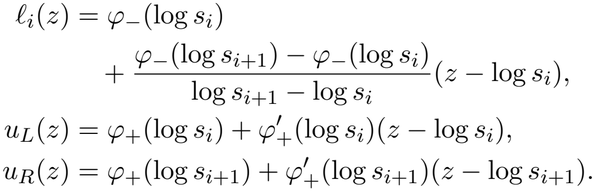

By concavity, we have that *ℓ*_*i*_ ≤ *φ*_*−*_ ≤ *φ*_+_ ≤ min {*u*_*L*_, *u*_*R*_}, which, with the bounds on *k*, gives us the desired inequality (5). We combine these rate bounds with the remaining equilibrium balances and obtain their projections, as in Theorem 4 to obtain *g*_*i*_. See Appendix A for the refinement claim.

*Remark*. The guarantees above provide an envelope that bounds the extremal responses. The lower and upper loga-rithmic rate bounds differ from their corresponding extremal logarithmic responses by at most *n*^2^ log^2^(*s*_*i*+1_*/s*_*i*_)*/*32.

## VI. Canonical Examples

### A. Regulatory Biological Circuits

#### 1) Single protein expression

Consider a simple protein expression example with a Hill propensity for the production and constant decay rate. A model of this system is 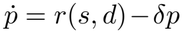, where *p* is the protein concentration, *s, d* are the regulator and DNA inputs, *δ >* 0 is the decay coefficient, and *r* is a proportional activation or repression propensity from Table 1. At equilibrium, *p*^*\**^ = *r*(*s, d*)*/δ*, so the contract guarantees follow from (5). A typical classification used by biologists is to divide the input into OFF, transition, and ON intervals. We apply this classification and impose the corresponding input bounds as assumptions to obtain the contracts visualized in Figure 1. The refined contracts give stronger equilibrium guarantees under the same assumptions, with adjacent contracts sharing their boundary values.

**Fig. 1.**
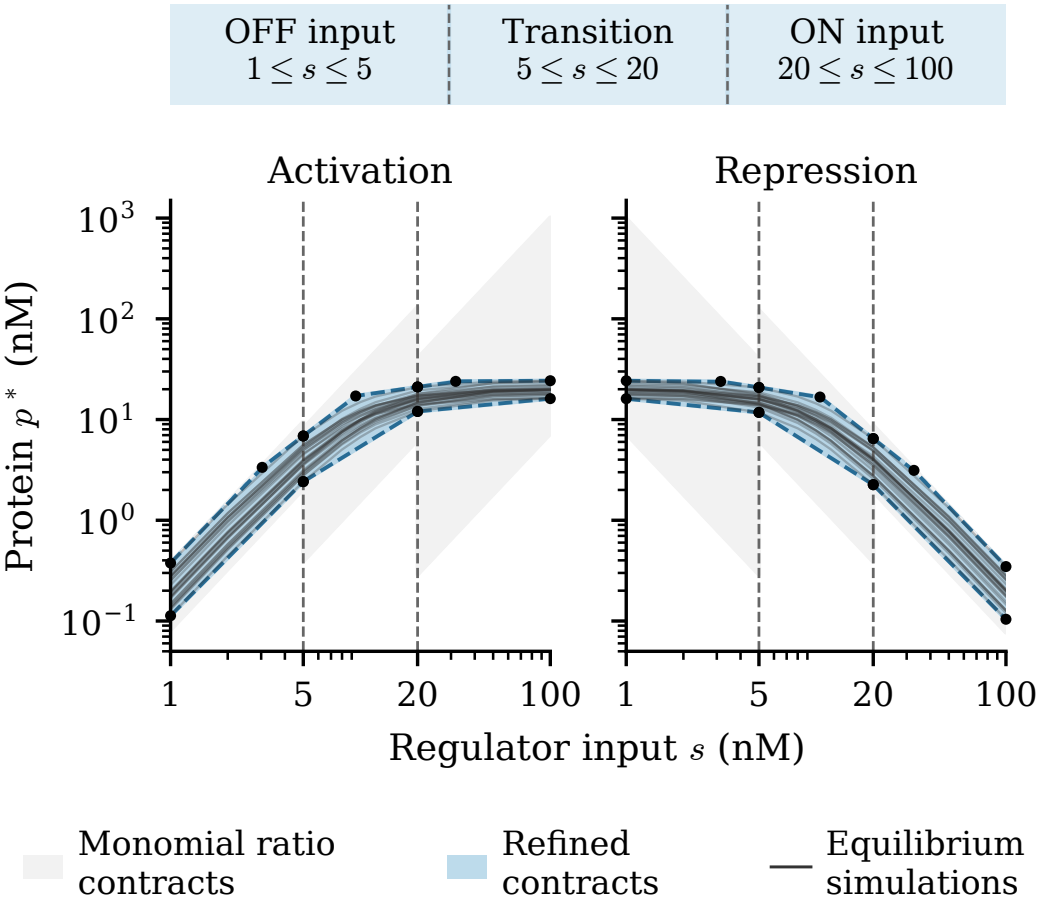
Equilibrium contracts for activation and repression with *n* = 2, *d* = 2 nM, *k ∈* [0.9, 1.1] min^*−*1^, *K ∈* [8, 12] nM, and *δ ∈* [0.09, 0.11] min^*−*1^. The contract assumptions are annotated.

#### 2) Composition of protein expression subsystems

Now we consider the design of a regulated protein expression system where a protein *p*_1_ represses the transcript of *p*_2_, the protein of interest. We specify this design with a top-level requirement: 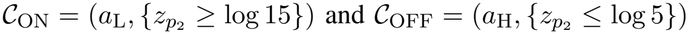,where *a*_L_, *a*_H_ are DNA input assumptions for low and high conditions, respectively. To meet these requirements, we model two transcription and translation modules with DNA inputs *d*_1_, *d*_2_. Each module has dynamics 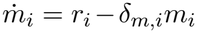 and 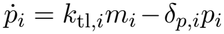 where *m*_*i*_, *p*_*i*_ are transcript and protein concentrations, *k*_tl,*i*_ is the translation coefficient, and *δ*_*m,i*_, *δ*_*p,i*_ *>* 0 are degradation coefficients. The transcription rates are *r*_1_ = *k*_1_*d*_1_, with *k*_1_ *>* 0, and *r*_2_ = *r*(*p*_1_, *d*_2_), a proportional negative Hill propensity. At equilibrium, 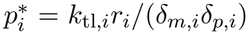 We derive the constitutive expression contract *C*_1_ from this balance and the repression contract *C*_2_ using Theorem 5.

With *d*_2_ = 1 nM, we choose the DNA input regions *a*_L_, *a*_T_, *a*_H_ so that the guaranteed range of 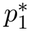 lies within [1, 5], [5, 20], and [20, 100] nM, respectively. Using Pacti [19], we apply `compose` to compute *C*_1_ *∥ C*_2_ that eliminates the shared variable *p*_1_ and gives us a guarantee on *p*_2_. Then, we use `optimize` to check the refinement inequalities by linear programming. We get that the composed contracts refine *C*_ON_ and *C*_OFF_ under their respective assumptions, as visualized in Figure 2.

**Fig. 2.**
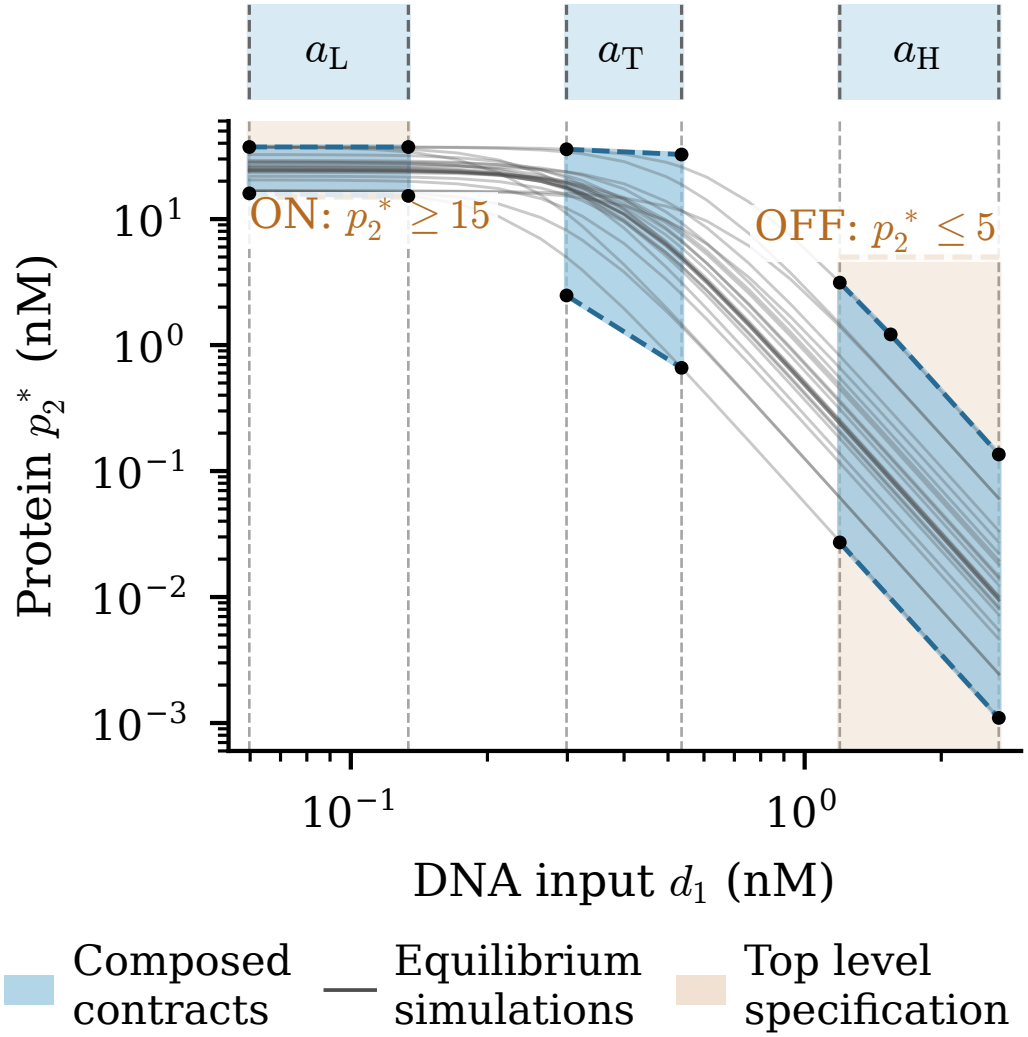
Composed guarantees under the DNA input assumptions *a*_L_, *a*_T_, *a*_H_ with *d*_2_ = 1 nM, *n* = 4, *K ∈* [9, 11] nM. The top level specification requires 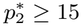 nM under *a*_L_ and 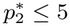 nM under *a*_H_. The *•* denotes polyhedral bound endpoints.

*Remark*. For the single protein activation example at *s* = 100 nM and *d* = 2 nM, the monomial ratio contract from Theorem 4 gives 7.23 ≤ *p*^*\**^ ≤ 1011.03 nM, which cannot certify the top level requirement *p*^*\**^ ≥ 15 nM. On the other hand, the refined contract gives 16.13 ≤ *p*^*\**^ ≤ 24.29 nM and certifies this requirement. Thus, abstraction refinement allows us to strengthen component guarantees under the same assumptions and certify requirements that the broader contracts could not. The stronger component contracts can then be substituted in compatible compositions, allowing these improvements to propagate to guarantees for the overall circuit.

### B. Formal Specifications for Enzymatic Output Flux

Next, we consider an enzymatic CRN subsystem:

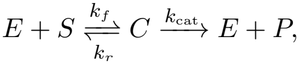

where *E, S, C, P* denote the enzyme, substrate, complex, and product concentrations, and *k*_*f*_, *k*_*r*_, *k*_cat_ *>* 0 are rate constants. We consider the complex balance at fixed enzyme and substrate totals, with conservation constraints *E* + *C* = *E*_tot_ and *S* + *C* = *S*_tot_. We set *x* = *S*_tot_, *u* = *E*_tot_, and 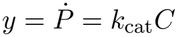.

Using monomial dominance, we get the logarithmic derivative guarantees from Theorems 2 and 3. We show the full derivation in Appendix B. The sensitivity polytope, conv { (1, 1), (1, 0), (0, 1) }, lies in the nonnegative orthant. From Lemma 3, the output flux is therefore nondecreasing in both *x* and *u*. We use *z*_*x*_ = log *x, z*_*u*_ = log *u, z*_*y*_ = log *y, κ* = log *K*, and *κ*_cat_ = log *k*_cat_, where *K* = (*k*_*r*_ + *k*_cat_)*/k*_*f*_ . For the regime dominated by *K*, the contract is *C*_*K*_ = (*A*_*K*_, *G*_*K*_), with positive margins *δ*_*Ku*_, *δ*_*Kx*_ and

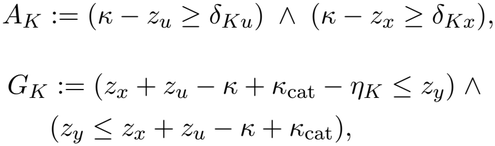

where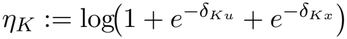 .

For the regime dominated by substrate, the contract is

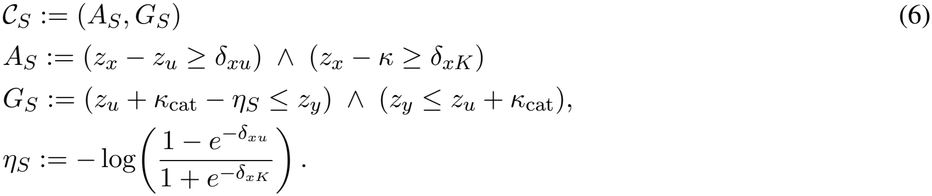

Finally, the regime dominated by enzyme, the contract *C*_*E*_ = (*A*_*E*_, *G*_*E*_) follows from (6) by interchanging *x* and *u*, with margins *δ*_*ux*_, *δ*_*uK*_ in place of *δ*_*xu*_, *δ*_*xK*_. Under the substrate excess assumption *S* ≈ *x*, the Michaelis Menten response is *y*_MM_ = *k*_cat_*ux/*(*K* + *x*). Using Theorem 5 with *n* = 1, *s* = *x, d* = *u*, and *k* = *k*_cat_, we obtain the contract *C*_MM_ = (*A*_MM_, *G*_MM_) under the assumptions *z*_*x*_ *z*_*u*_ ≥ *δ*_MM_ and *z*_*x*,low_ ≤ *z*_*x*_ ≤ *z*_*x*,high_. These contracts bound the output flux under explicit enzyme and substrate conditions (Figure 3).

**Fig. 3.**
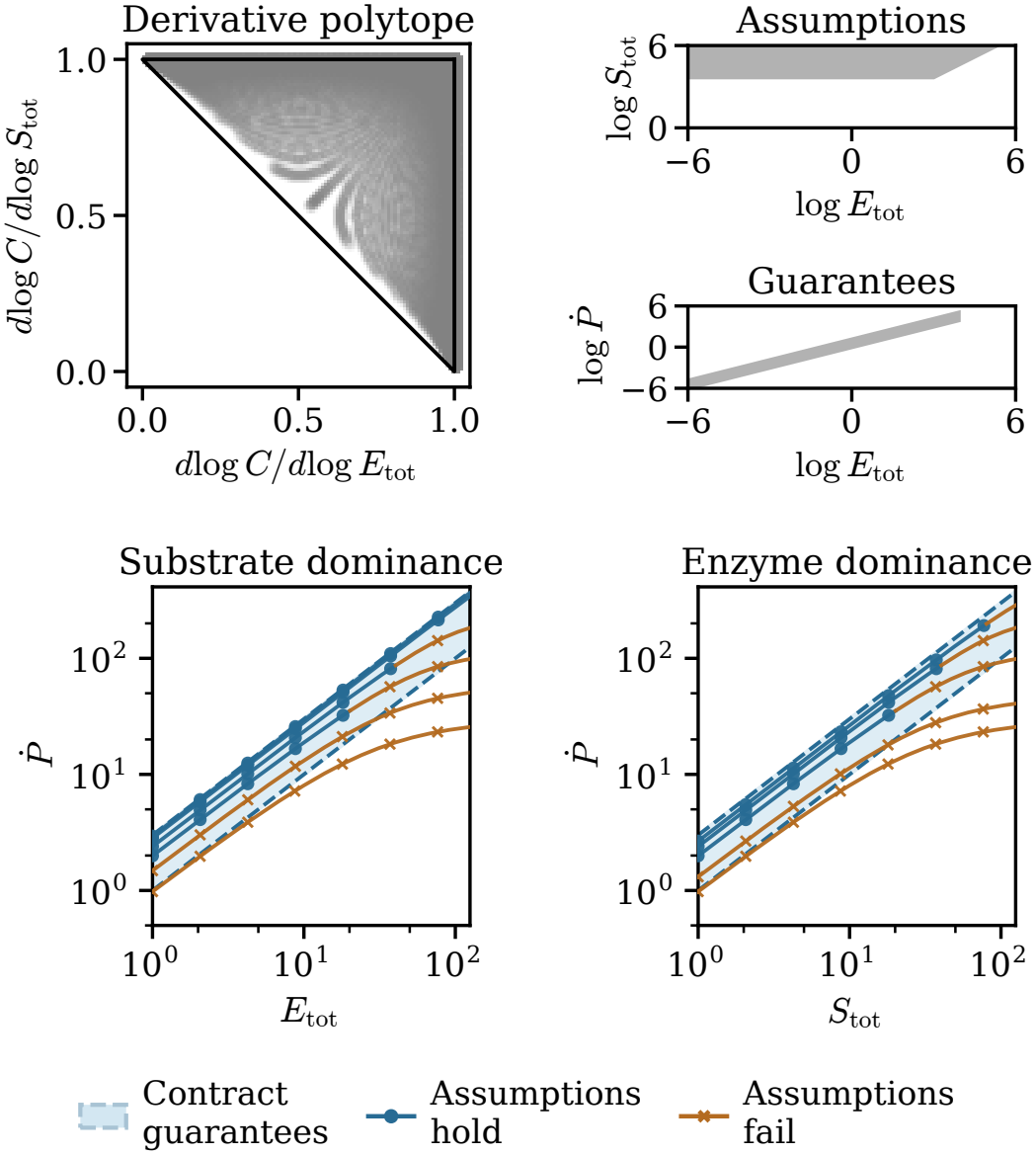
Logarithmic derivative guarantees and output flux contracts for the enzymatic subsystem. We observe that, for *S*_tot_ = 40, the substrate dominance assumption fails when *E*_tot_ *>* 20, and the equilibrium flux falls below the contract bound 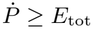 when *E*_tot_ *>* 90.

## VII. Biocircuit Design Applications

In this section, we consider three biocircuit design applications that have experimental basis from the literature: (1) constructing a library of logic gates from Cello [7], (2) mechanistic design space exploration to achieve a strict logic gate specification [26], and (3) selecting the RBS and promoters of a circuit by screening a library of components.

For each subsystem (Σ_*i*_) CRN, the theoretical results we have presented can be used to write a contract C_*i*_. Then, using the standard contract composition rule [19], [23], the interconnected system has *C*_full_ = *C*_1_ *∥* · · · *∥ C*_*N*_ . Given a desired system level contract *C*_spec_, we address the compositional design and analysis problem by checking *C*_full_ *⪯ C*_spec_.

### A. Library of logic gate contracts using Cello

Cello is a biocircuit design tool that uses a characterized genetic logic gates library to compose parts that meet a specification in Verilog [7]. Specifically, the Cello gate library provides characterizations for 20 NOR logic gates and 2 AND logic gates in *E. coli*. We use these characterizations to construct a library of polyhedral assume guarantee contract specifications for compositional design with the same characterized biological parts. Our constructed library of contracts then enables designers to express top level requirements beyond just logical requirements in Verilog. For example, one may prescribe admissible input ranges, required output levels, and other quantitative relations between inputs and outputs in a top-level contract and then reason about what parts from Cello meet these requirements.

For the NOR gates in Cello, the response function is

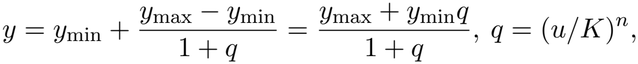

where *u* is the aggregate input promoter activity and *y* is the output promoter activity, both in RPU. For the AND gates, the two input promoter activities *u*_1_, *u*_2_ give

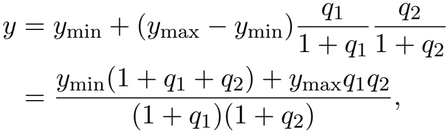

where *q*_*i*_ = (*u*_*i*_*/K*_*i*_)^*n*^ for *i ∈* {1, 2}. The Cello library parameters satisfy 0 *<y*_min_ *<y*_max_, with positive thresholds *K, K*_*i*_ and exponent *n*. Both response functions have the positive monomial sum quotient form in Definition 2. We write each as *y* = *A*(*u*)*/B*(*u*) and apply Theorem 4 to the balance *yB*(*u*) = *A*(*u*), then refine its guarantees using Proposition 2.

We divide each input into low, transition, and high intervals and construct a contract for each of the three joint regions. The assumptions are given by the following conditions on input *i*:

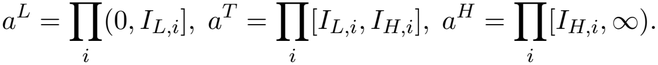

where *I*_*L,i*_ *<I*_*H,i*_ are the low and high input thresholds. For a NOR response, the aggregate input is *u* = *u*_1_ + *u*_2_. For an AND response, each assumption places conditions on both *u*_1_ and *u*_2_. We obtain three contracts for each characterized gate, giving a library of 66 contracts for the 22 response curves. These contracts provide formal component specifications for compositional design with the characterized parts in Cello. We illustrate one NOR contract using P1 PhlF visually in Figure 4, with the monomial ratio-derived guarantees and the tighter refined guarantees shown.

**Fig. 4.**
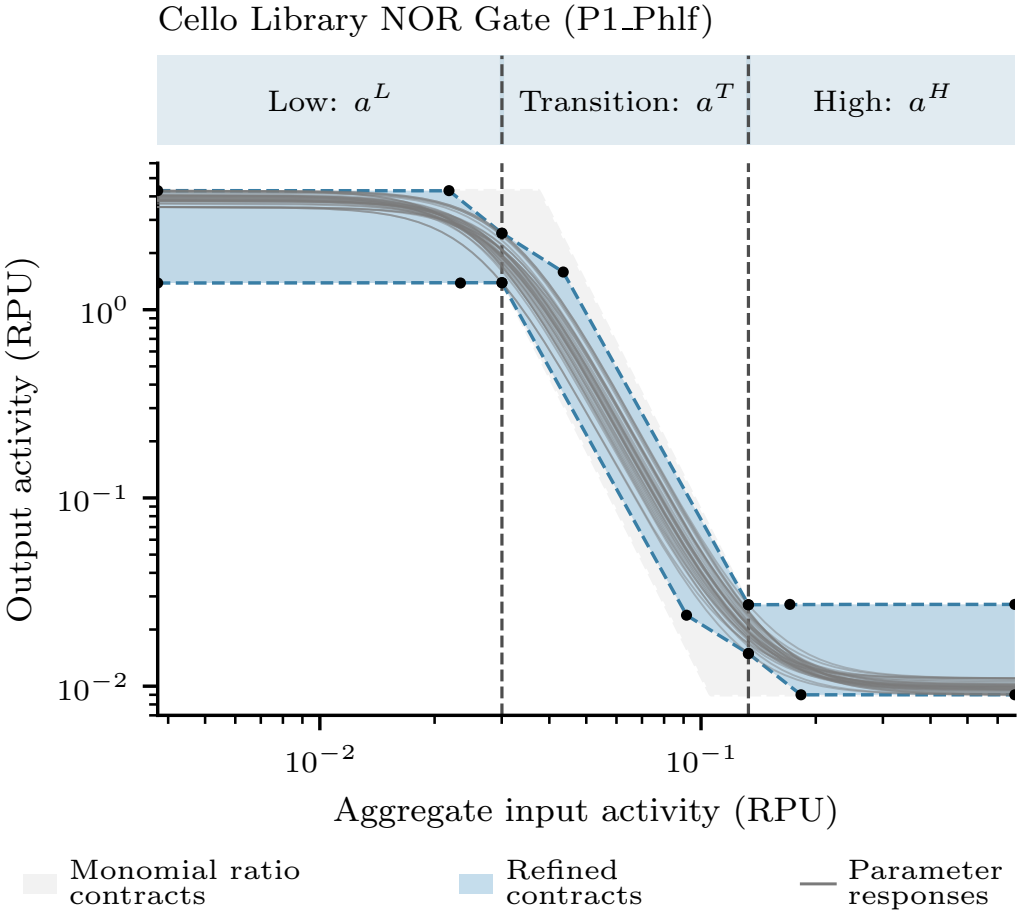
Input assumptions and refined output guarantees for P1_PhlF from the Cello gate library. The *•* denotes polyhedral bound endpoints.

### B. Regulation of Output Flux in an AND Logic Gate

We consider the design of a genetic circuit that implements a two-input AND logic circuit.

#### 1) Design goal

We express our requirement for the AND logic gate using the following top-level specification contract on inputs *ξ*_1_, *ξ*_2_ and output flux *y*:

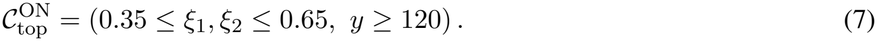

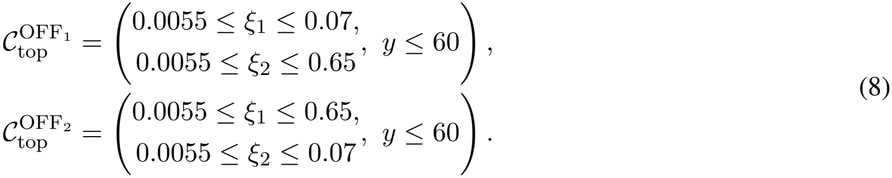

The input thresholds here are based on published data for promoter activity ranges in the Marionette library [2] and the characterized leak floors in Table 2. The maximum OFF input activity, 0.07 RPU, is chosen to be 20% of the minimum ON input activity, 0.35 RPU, and we do not require any Boolean behavior in the transition interval 0.07 *<ξ*_*i*_ < 0.35. The requirement on the output flux of gene expression is prescribed using common kinetic parameters and to ensure a twofold separation between the largest allowed OFF flux and the smallest allowed ON flux. The three operating modes of the circuit define a contract family *ℭ*_spec_ = {*C*_ON_, *C*_OFF,1_, *C*_OFF,2_ }. We say that a design *D* satisfies the complete specification when

**TABLE II.** Sensor parameters that guide *C*_spec_.

| Sensor | Promoter | Regulator | Leak | Hill activation | Cooperativity |
|---|---|---|---|---|---|
| aTc | *P*<sub>Tet\*</sub> | TetR | 0.0049 | 0.013 | 3.8 |
| Cuma | *P*<sub>CymRC</sub> | CymR<sup>AM</sup> | 0.0043 | 8.9 | 2.4 |

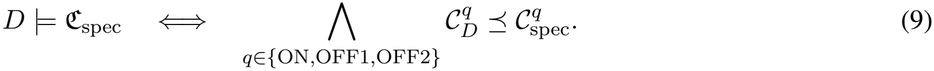

Thus, for each mode, refinement preserves the intended conditional guarantees. In defining the top-level specification, we have considered that for design proposals, a physical inducer concentration can be mapped to the promoter activity *ξ*_*i*_. This common RPU interface lets a sensor be connected to either expression subsystem. Therefore, regardless of the sensor identity and the assignment order, the downstream contracts variables become identical after interface abstraction.

#### 2) Candidate genetic circuits and interface contracts

We consider three possible mechanisms to achieve the AND gate specification: the HrpR/HrpS *σ*^54^ joint activation circuit [26], split T7 RNA polymerase complementation [27], and the InvF/SicA activator and chaperone gate [28]. In each circuit, two input promoters drive expression of components *Y*_1_ and *Y*_2_, which associate to form an activating complex *C*_3_ that drives reporter production. We assume that binding equilibrates faster than protein expression. Let *ξ*_*i*_ be promoter activity in RPU, *T*_*i*_ the total concentration of component *i* in nM, and *g*_*i*_ the expression gain in nM/RPU. At equilibrium, *T*_*i*_ = *g*_*i*_*ξ*_*i*_. The dissociation constant is *K* = *k*_*r*_*/k*_*f*_, where *k*_*f*_ and *k*_*r*_ are the association and dissociation rate constants. The reporter production rate constant is *k*_out_ in min^*−*1^, and the output flux is *y* = *k*_out_*C*_3_ in nM/min. The component identities and nominal parameters are given in Table 3. See Appendix C for the full model and equilibrium derivation.

**TABLE III.** Candidate AND gate designs and nominal parameters.

| Component | HrpR/HrpS [26] | split T7 [27] | InvF/SicA [28] |
|---|---|---|---|
| Subsystem 1 | HrpR (*Y*<sub>1</sub>) | Amino fragment (*Y*<sub>1</sub>) | InvF (*Y*<sub>1</sub>) |
| Subsystem 2 | HrpS (*Y*<sub>2</sub>) | Carboxyl fragment (*Y*<sub>2</sub>) | SicA (*Y*<sub>2</sub>) |
| Complex | HrpR/HrpS (*C*<sub>3</sub>) | Polymerase (*C*<sub>3</sub>) | InvF/SicA (*C*<sub>3</sub>) |
| Reporter | *P* | *P* | *P* |
| *K* | 250 | 1200 | 250 |
| *k*<sub>out</sub> | 1.2 | 10 | 1.0 |
| *g*<sub>1</sub> | 4000 | 500 | 800 |
| *g*<sub>2</sub> | 800 | 700 | 4000 |

For the mode *q ∈* {ON, OFF1, OFF2}, we define 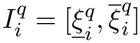 as its promoter activity interval. For any candidate circuit *d*, we have the following assumptions

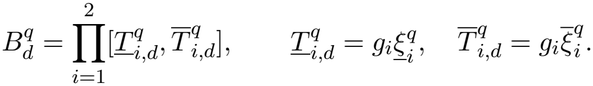

We bound the equilibrium reporter flux *f*_*d*_(*T* ) = *k*_out_*C*_3_(*T* ; *K*) over 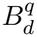 at the selected nominal parameters. The flux is nondecreasing in both component total concentrations, *T*_1_ and *T*_2_ (Appendix C), so its lower and upper bounds occur at the corresponding concentration endpoints: 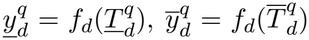. The subsystem contracts are therefore

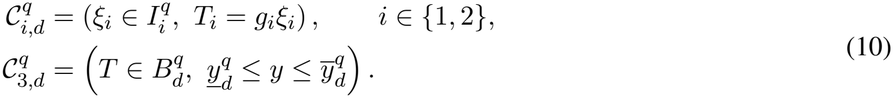

We compare the three circuits against the top-level ON and OFF requirements using these contracts.

#### 3) Compositional verification

For every mode, the expression guarantees in (10) establish the assumption 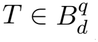 of 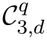 . The subsystem contracts compose as 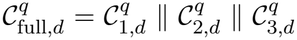. We obtain the external guarantee by projecting out the internal interface quantities. To refine the specification, the composed contract must admit every required external input, and every permitted output under those inputs must meet the corresponding ON or OFF requirement. Thus refinement of (7) and (8) requires

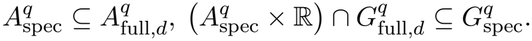

The complete specification (9) is satisfied only when these conditions hold for every mode. We compare the three candidate circuits with both sensor assignments and record the ON and OFF verification decisions in Table 4.

**TABLE IV.**
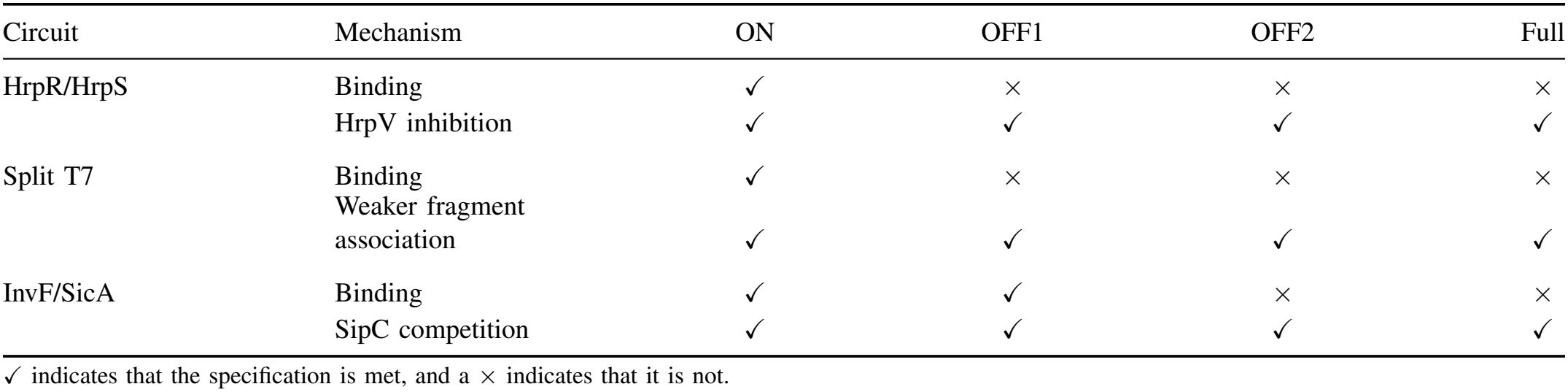
Compositional verification of the AND specification.

#### 4) Verification results and circuit redesign

At the nominal parameter choices, none of the three binding circuits satisfies the complete AND specification. All three contracts pass the ON check but HrpR/HrpS and split T7 fail both OFF checks, while InvF/SicA passes OFF1 and fails OFF2. We therefore propose changes to the circuit designs. We select the HrpV concentration, the strength of association between the split T7 fragments, and the SipC production flux as the target design changes. These choices are made since these mechanisms are involved in suppressing the OFF output while preserving the required ON output.

For HrpR/HrpS, HrpV inhibits reporter production by binding the activating complex. Under fast and strong inhibitor binding, reporter production begins when the complex concentration exceeds the inhibitor total *V*_tot_. We select *V*_tot_ = 120 nM to suppress the OFF output while allowing sufficient ON output. The resulting reporter relation is given in Appendix C.

For split T7, the amino and carboxyl polymerase fragments must associate to restore transcriptional activity. We consider weaker fragment association, represented by a larger dissociation constant *K*. We select *K* = 3000 nM so that the ON output remains sufficient while both OFF outputs decrease. We construct the circuit contract using the same binding balance and the bounds in Appendix C.

For InvF/SicA, SipC competes with InvF for binding to SicA. Production, degradation, SipC secretion, and competition for SicA determine the concentration of the transcriptionally active InvF/SicA complex. The input expression contracts determine production fluxes *p*_*i*_ = *γ*_*i*_*T*_*i*_ for *i ∈* {1, 2}, where *γ*_*i*_ is the degradation rate constant of component *i*. We select a SipC production flux *p*_*Z*_ = 149.45 nM/min and construct the reporter flux contract in (10). The full reaction dynamics, existence and uniqueness of the equilibrium, and monotonicity of reporter flux in the two input production fluxes are provided in Appendix C.

We show that the three contract analysis-driven circuit redesigns satisfy the ON and OFF requirements of the AND specification (see bottom panel in Figure 5). A limitation of this application example is that we did not account for resource loading or other feedback in circuits, which are known to play an important role in circuit performance [11], [29].

**Fig. 5.**
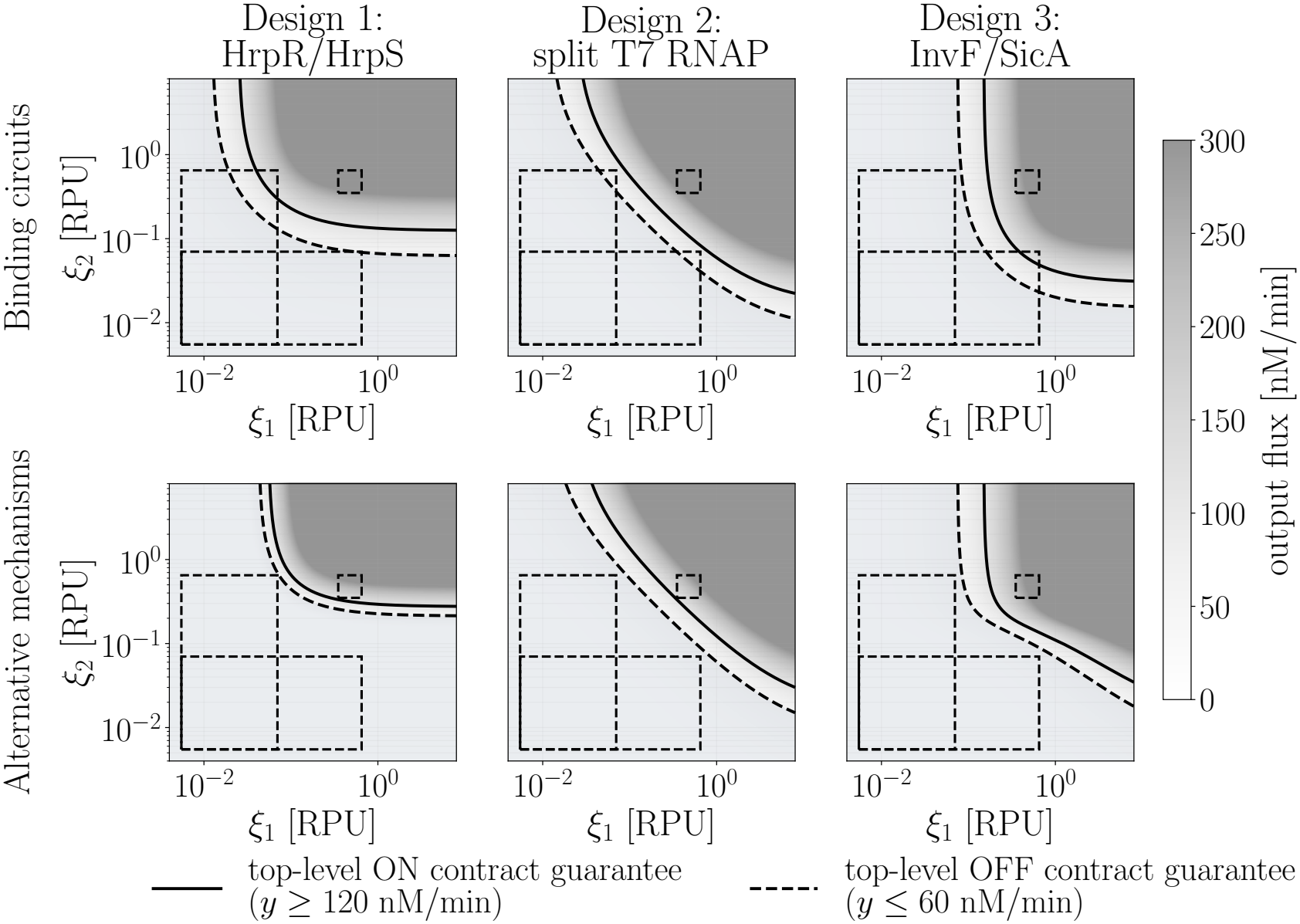
Verification of the AND specification for the binding circuits fails for the initial design. The proposed redesign mechanisms at the selected nominal parameter values (bottom row) meet the desired specification.

### C. RBS Library Screening for Biocircuit Design

Wetlab scientists designing biomolecular circuits often need to select components from characterized libraries, such as the CIDAR MoClo kit and libraries of promoters and regulatory elements [2], [7], [30]. For this selection, an experiment needs to be run that screens through the library to choose the right components. In this application example, we show how component contracts can be used to select RBS assignments that satisfy a top-level NAND logic specification.

We consider the transcriptional NAND circuit in [26] where inducible promoters *P*_lac_ and *P*_BAD_ express HrpR and HrpS expression, respectively. Then, these two regulators jointly activate *P*_hrpL_, which leads to the expression of the lambda repressor CI. Since CI represses the reporter, the fluorescence output exhibits a NAND logic behavior.

We denote the HrpR and HrpS expression levels by *R* and *S*, the CI level by *C*, and reporter fluorescence by *G*. The key design choices are to select three RBSs for the circuit, one for each of the two input promoters and one for tuning the CI expression gain *g*.

To parameterize the component models, we use data from [26] (promoter characterizations in Table 2, the AND characterization in Supplementary Figure S6, and the rbs34 repression characterization in Supplementary Table S1 of [26]). This data is for *E. coli* MC1061 in M9 glycerol medium at 30 ^*°*^C. We label the reported working circuit in their paper as the ground truth goal for our design exploration of RBS library screening. This circuit uses rbsH with *P*_lac_ and rbs33 with *P*_BAD_.

#### 1) Top-level specification

With two inputs, *I*_1_ and *I*_2_ (IPTG and arabinose in mM), we require a top-level specification on the normalized fluorescence output, *F* = *G/k*_*G*_, where *k*_*G*_ is the calibrated reporter amplitude. Our goal is to find designs that satisfy the top level contract family *C*_top_ = {(*a*_*q*_, *g*_*q*_) : *q ∈* {OFF, ON1, ON2}}, with

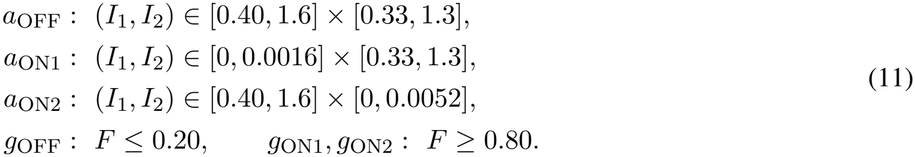

#### 2) Equilibrium Component Interfaces

Regulator binding activates expression in the two input modules. The AND promoter requires occupancy by both HrpR and HrpS to produce CI, and CI binding represses reporter transcription. The promoter dynamics, conservation laws, and equilibrium derivations are given in Appendix D. For independent binding of the two regulators, the equilibrium active AND promoter fraction *a* and CI expression level satisfy

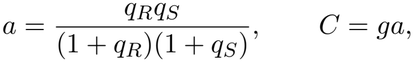

where *q*_*R*_, *q*_*S*_ are normalized regulatory activities, and *g >* 0 is the CI expression gain. The unoccupied reporter promoter fraction *w* gives the normalized fluorescence

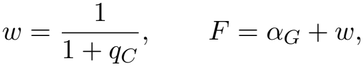

where *q*_*C*_ is the CI regulatory activity and *α*_*G*_ is the dimensionless basal transcription coefficient.

We relate inducer concentrations and expression levels to regulatory activities using the fitted relations in

Supplementary equations S2, S3, and S4 of [26]:

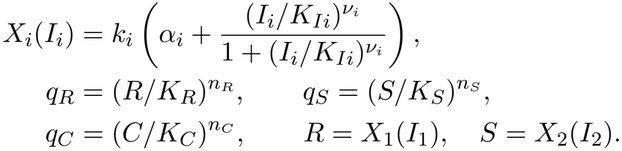

The parameters *k*_*i*_, *K*_*Ii*_, *ν*_*i*_, *α*_*i*_ are the input response amplitude, inducer threshold, Hill exponent, and basal coefficient, respectively (values in Table 5). The thresholds *K*_*R*_, *K*_*S*_, *K*_*C*_ and exponents *n*_*R*_, *n*_*S*_, *n*_*C*_ describe the AND and repression activity maps. We use dimensionless contract coordinates *u*_*i*_ = *I*_*i*_*/*(1 mM), *z*_*R*_ = log(*R/K*_*R*_), *z*_*S*_ = log(*S/K*_*S*_), *z*_*C*_ = log(*C/K*_*C*_), *z*_*a*_ = log *a*, and *z*_*w*_ = log *w*. By continuity of the interface quantities and closedness of the contract inequalities, we note that the guarantees will also extend to the boundary.

**TABLE V.**
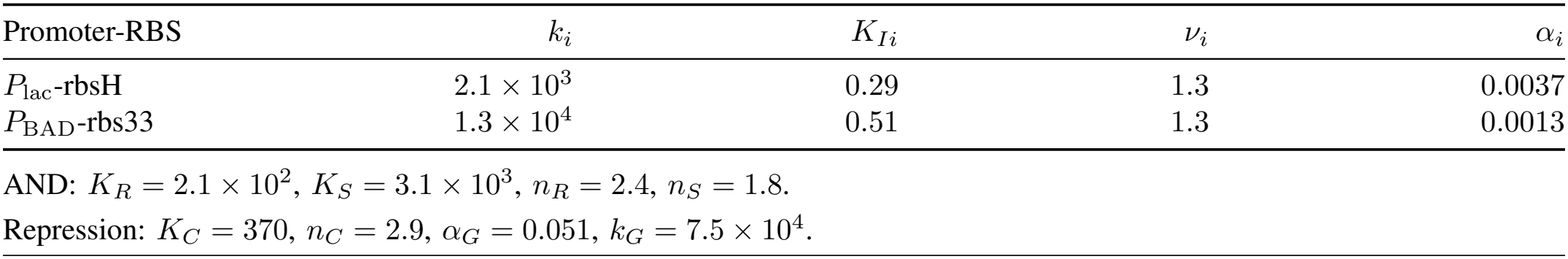
Circuit parameters for the reference RBS assignment.

| Promoter-RBS | *k*<sub>*i*</sub> | *K*<sub>*Ii*</sub> | ν<sub>*i*</sub> | α<sub>*i*</sub> |
|---|---|---|---|---|
| *P*<sub>lac</sub>-rbsH | 2.1 × 10<sup>3</sup> | 0.29 | 1.3 | 0.0037 |
| *P*<sub>BAD</sub>-rbs33 | 1.3 × 10<sup>4</sup> | 0.51 | 1.3 | 0.0013 |
| AND: *K*<sub>*R*</sub> = 2.1 × 10<sup>2</sup>, *K*<sub>*S*</sub> = 3.1 × 10<sup>3</sup>, *n*<sub>*R*</sub> = 2.4, *n*<sub>*S*</sub> = 1.8. |  |  |  |  |
| Repression: *K*<sub>*C*</sub> = 370, *n*<sub>*C*</sub> = 2.9, α<sub>*G*</sub> = 0.051, *k*<sub>*G*</sub> = 7.5 × 10<sup>4</sup>. |  |  |  |  |

#### 3) Contract construction and verification

We construct component contracts from the equilibrium maps using Theorem 4 and the positive sum bounds of Lemma 4 while Lemma 1 gives affine ratio assumptions in logarithmic coordinates. On each required input region, monotonicity of the calibrated promoter responses gives expression bounds 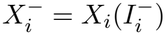 and 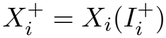. So we have the extremes

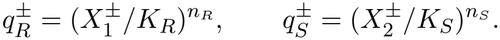

that set up the intervals for the input assumptions of the AND contract. The CI expression balance then gives the input bounds for the repression contract. Thus, each RBS assignment has component contracts constructed from its own calibrated expression ranges. We visualize the contracts for one reference RBS choice in Figure 6. The upstream contracts give an AND promoter activity interval 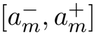 for each required input region *m* and the repression mechanism converts a fluorescence threshold *F*_*\**_ into the CI threshold

**Fig. 6.**
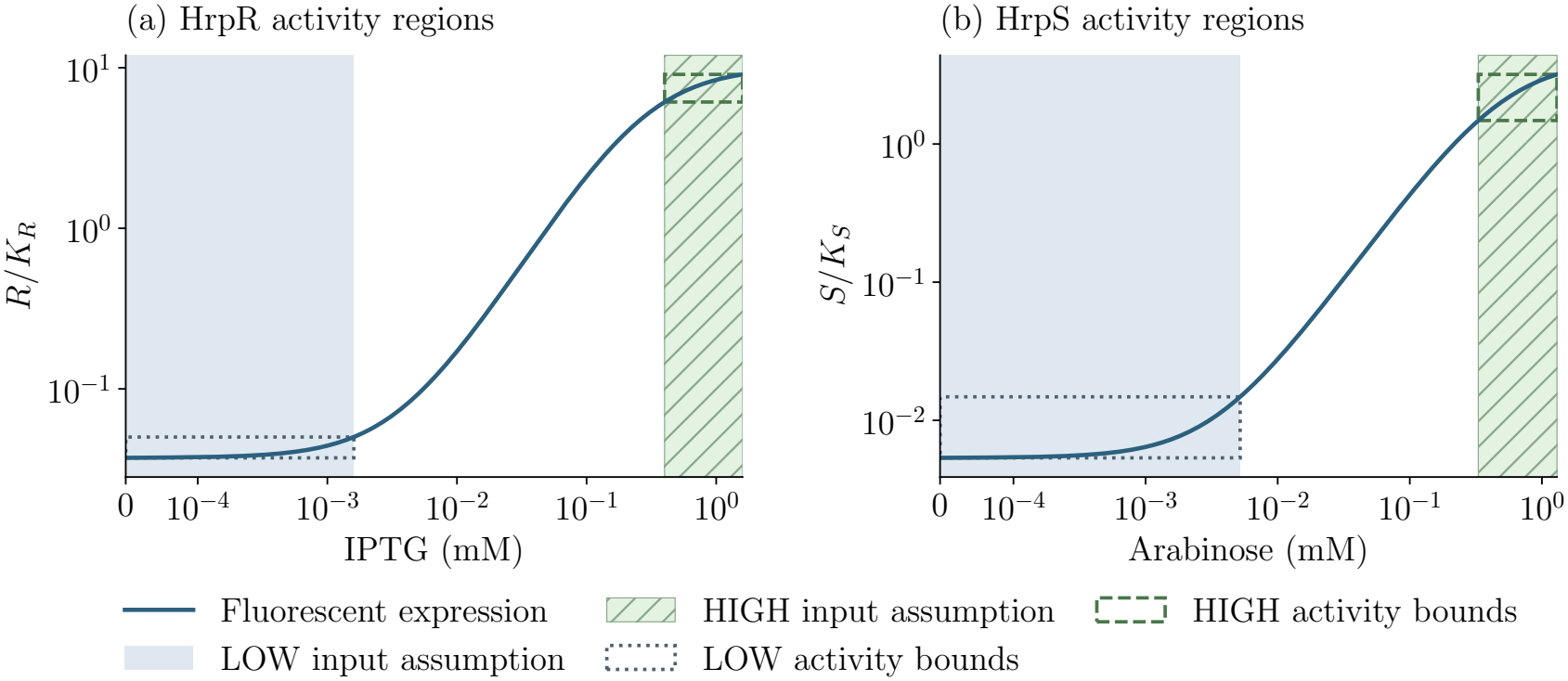
Inducer assumptions and calibrated HrpR and HrpS activity for the ground truth choice of RBS from [26].

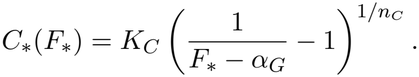

The OFF requirement gives a lower bound on *g*, and the ON1/2 requirements give upper bounds. A sufficient gain satisfies

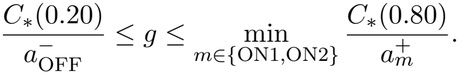

For the reference RBS choice, we select *g* = 1.8 × 10^4^, corresponding to *g/K*_*C*_ = 49. We compose the regulator expression, AND mechanism, and the repression component contracts at this gain using Pacti. The composed contracts satisfy the fluorescence requirements throughout all three input regions. The OFF output bound is below 0.051, and the ON1 and ON2 output bounds are above 1.0 under the amplitude normalization in (11). Also, we note that monotonicity of the equilibrium response gives the HIGH fluorescence condition when both inputs are LOW.

#### 4) Formal library screening

We construct and compose contracts using the calibrated promoter activity ranges for each RBS choice in the RBS library. With this contract-based library screening, we are able to certify seven RBS choices, establish violations for 16, and leave 7 choices unresolved. The library comprises 30 assignments from five usable *P*_lac_ calibrations and six *P*_BAD_ calibrations. We note that *P*_lac_/rbs33 was discussed in [26] as an unsuccessful choice because it showed no response to IPTG induction.

The 4 certified pairs that meet the required specification are *P*_BAD_/rbs33 with *P*_lac_/rbs30, rbs32, rbs34, or rbsH. The other three pairs are *P*_BAD_/rbs32 with *P*_lac_/rbs30, rbs34, or rbsH. Each of the 16 rejected assignments has a composed fluorescence bound that contradicts at least one requirement of the top level NAND contract family *C*_top_ in (11). In Figure 7, each row identifies the *P*_lac_ RBS and columns identify the *P*_BAD_ RBS. We compare the response curves with the composed guarantees over the eight published inducer doses (labeled on the X-axis).

**Fig. 7.**
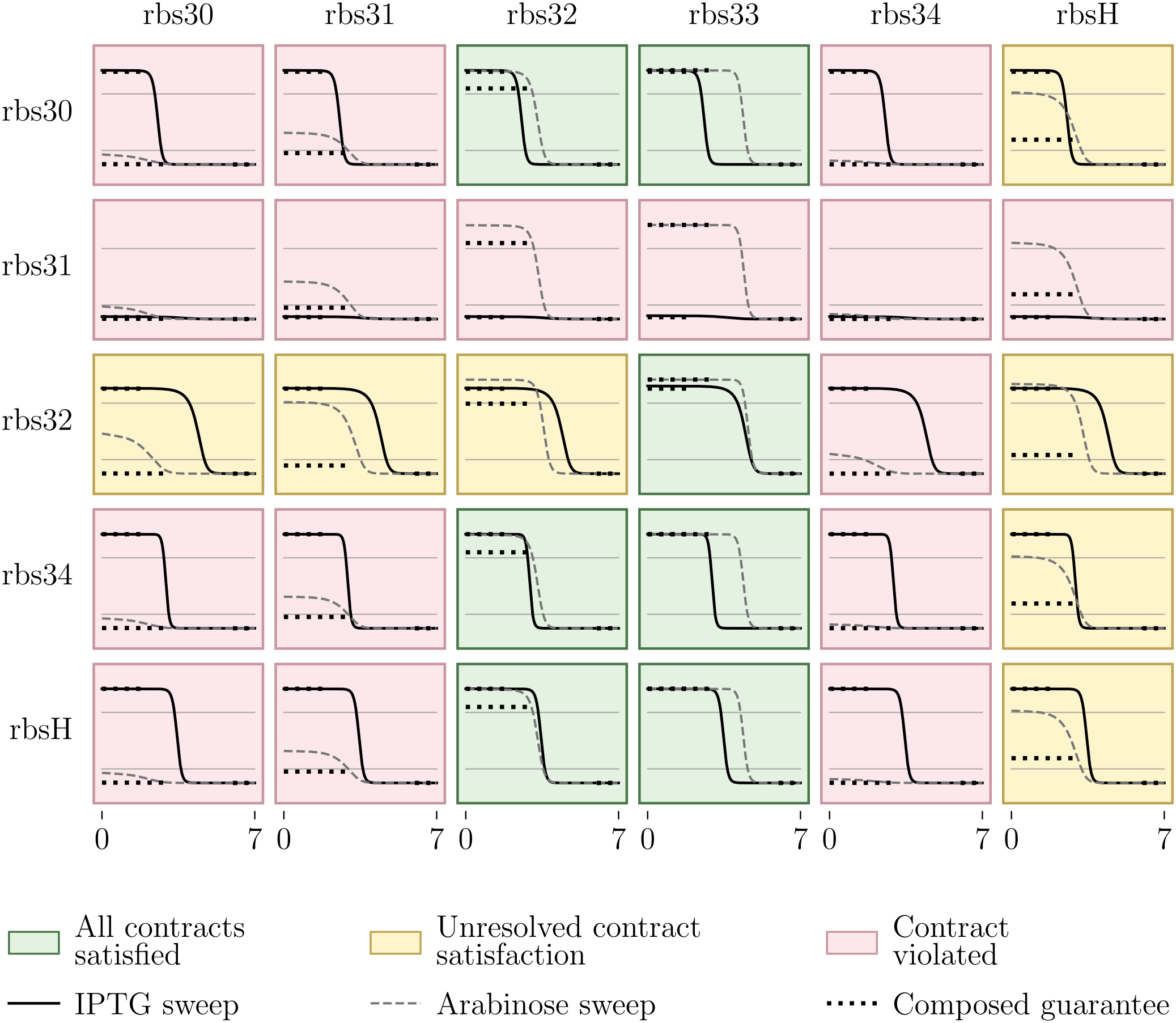
Formal screening of 30 RBS assignments using calibrated promoter activity ranges and composed fluorescence guarantees.

#### 5) Comparison with published data

We compare the calibrated equilibrium model response and the composed contract bounds for the reference RBS assignment with the published response of the assembled NAND circuit. We use the mean fluorescence values from Figure 5(c) of [26]. We normalize the model predictions and contract bounds for *F* = *G/k*_*G*_ in (11) by its maximum model value of 1.05 to compare relative fluorescence with the published circuit response.

The largest prediction errors occur at low IPTG and high arabinose with MAE of 0.361 in ON-1, 0.032 in ON-2, and 0.037 in OFF (see Figure 8). This discrepancy is expected due to the leaky HrpR expression, discussed in [26]. Across all 28 experimental observations, we find a high Pearson correlation (0.893) between predicted and measured relative fluorescence. Twelve of the experimental observations are clearly within the required input regions, and six of the refined contract bounds are within the experimental error bars shown in Figure 8. So, the predictions agree closely with the measured high fluorescence in ON-2 and low fluorescence in the OFF condition (the primary NAND output).

**Fig. 8.**
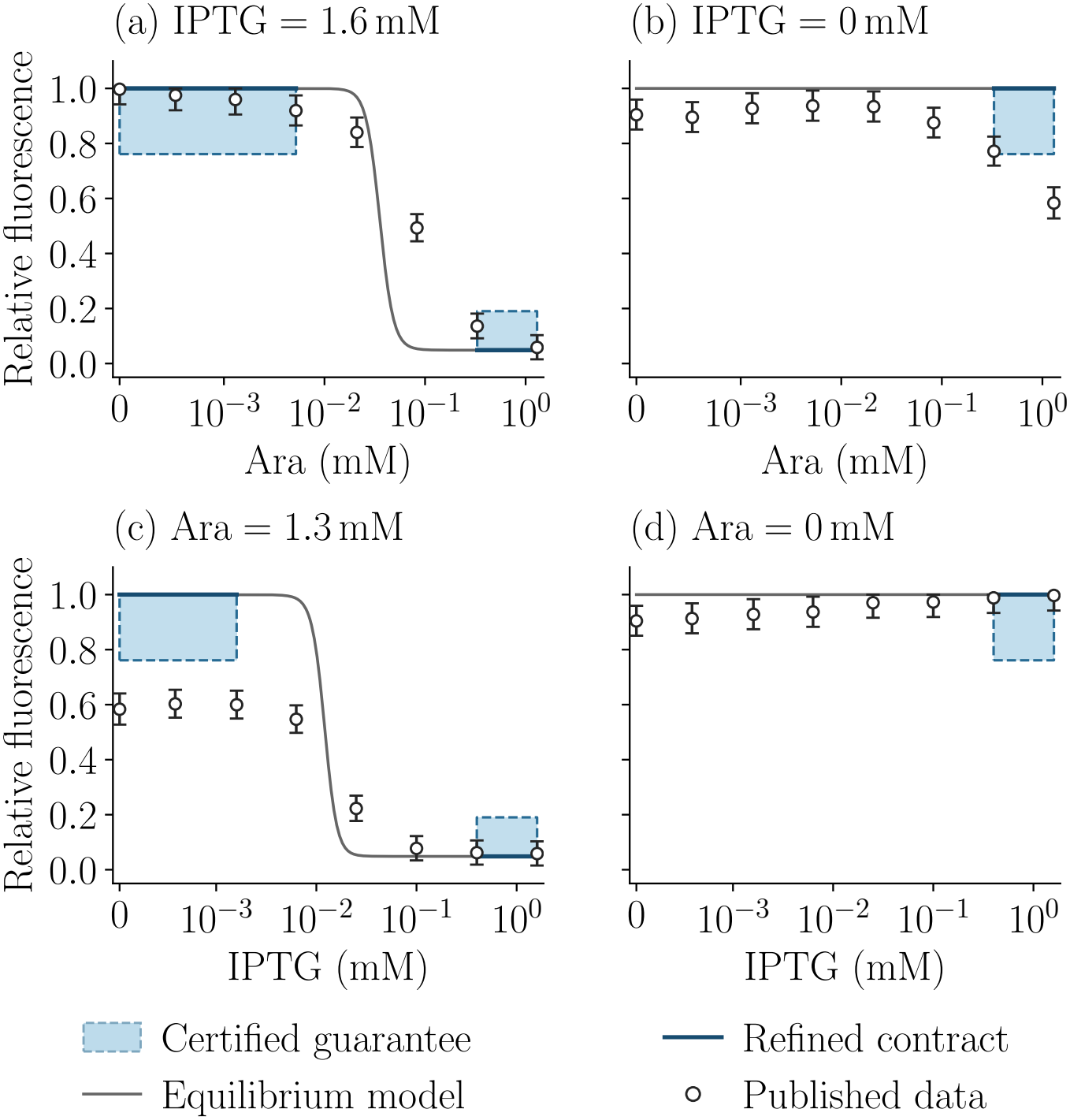
Equilibrium model response and contract guarantee regions compared with published data [26] along four boundary sweeps. The shaded regions show the certified OFF and ON guarantees in (11) with physical output bounds, and the narrow bands show the refined contracts. The error bars report the error in scraping the values from the graph images published in the paper.

## VIII. Conclusion

We present a new method for deriving assume guarantee contracts for biomolecular systems from their CRN descriptions using exponent polytopes and bounds on positive monomial sums. This establishes a theoretical basis for expressing common modeling conditions for biomolecular systems: monomial dominance, conservation laws, and equilibrium balances. We show how CRNs under these conditions can be abstracted as formal contract specifications. Specifically, we derive contracts for mass action CRNs, equilibrium contracts for positive quotient CRNs, and for CRNs with standard Hill propensities. These contracts provide us with guarantees at every positive equilibrium under explicit operating assumptions. We illustrate our method using canonical biomolecular systems and synthetic biological circuit designs based on published experimental data. The canonical examples include enzymatic reactions, transcription and translation models for protein expression, and compositions of these systems. We find that the contract-based design approach makes subsystem operating assumptions explicit and enables composition and refinement checks against a top level system requirement. We show three different biocircuit design applications. Firstly, we show how to select biological mechanisms and parameter choices that realize an AND gate specification. Then, we show how we can construct a library of formal component specifications from the Cello gate library for compositional circuit design with characterized biological parts. Finally, we show how formal screening through contract composition, refinement, and certified output bounds can accelerate laboratory circuit design by rejecting incompatible RBS assignments and identifying choices that satisfy the required circuit specification. The accuracy of the predictions with published experimental data establishes the utility of our approach for compositional biocircuit design.

## Appendix A

### Refinement of the Hill contracts

#### Proof of Theorem 5 (continued)

To establish refinement, we subtract *τz*_*d*_ from the rate bounds in (5) and the corresponding monomial ratio bounds. Every reference affine lower bound is below log *k*^*−*^ + *φ*_*−*_ at both endpoints. But the new lower bound log *k*^*−*^ + *ℓ*_*i*_ equals log *k*^*−*^ + *φ*_*−*_ at these endpoints. So, the new lower bound is above the reference bound throughout the interval, because their difference is affine and nonnegative at both endpoints.

For the upper bounds, the reference slopes are 0 or *ν* = *n*(2*σ* 1), while 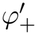 lies between these values. Then, a reference upper bound with the larger slope is above log *k*^+^ + *u*_*L*_ throughout the interval, and a reference upper bound with the smaller slope is above log *k*^+^ + *u*_*R*_. These comparisons follow from their endpoint values and the ordering of their slopes. Thus, the minimum of the endpoint tangents is below every reference upper bound. We then add the remaining equilibrium inequalities and project onto the interface variables. These operations preserve inclusion, which establishes 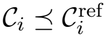 . Finally, at *z*_*s*_ = log *s*_*i*+1_, both neighboring lower chords equal *φ*_*−*_(log *s*_*i*+1_). Each neighboring pair of upper tangents contains the tangent at that endpoint, so its minimum equals *φ*_+_(log *s*_*i*+1_). The neighboring rate bounds therefore agree for every common value of *d*.

## Appendix B

### Enzymatic Output Flux

With the species and rates defined in Section VI-B, the complex dynamics and product flux are

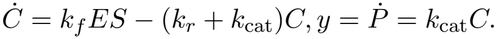

With *T*_1_ = *u, T*_2_ = *x, C*_3_ = *C*, and *K* = (*k*_*r*_ + *k*_cat_)*/k*_*f*_, the equilibrium satisfies the binding balance in Appendix C-A. This gives us the same result as in [18]:

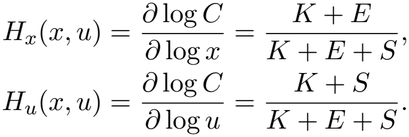

We can then write the convex decomposition in a two dimensional space of (*x, u*), that is, *H*(*x, u*) is

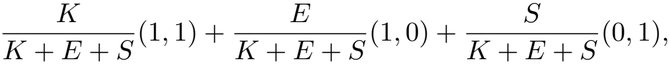

so *H*(*x, u*) *∈* conv {(1, 1), (1, 0), (0, 1)}, as shown in Figure 3. The three vertices correspond to the regime dominated by *K* (1,1), the regime dominated by enzyme (1,0), and a regime dominated by substrate (0,1), respectively, in the way these are conventionally described in the literature.

We now use the same equilibrium balance to derive the flux guarantees under the dominance assumptions. For the contract *C*_*K*_, the assumption *A*_*K*_ gives 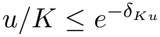 and 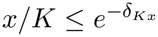. Then, (12) with *T*_1_ = *u* and *T*_2_ = *x* gives

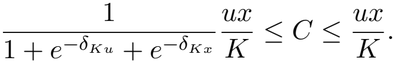

We multiply by *k*_cat_ and take logarithms to obtain *G*_*K*_. The substrate and enzyme dominance contracts use the

same equilibrium balance with the respective dominance assumptions.

## Appendix C

AND Circuit Models and Contract Derivations

We derive the equilibrium relations and monotonicity properties of the AND circuit models to obtain the output bounds in (10).

### A. Binding circuits

The two expression subsystems produce components that associate to form the reporter activating complex. The dynamics of the mass action CRN is given below

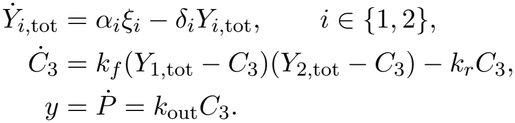

Here *Y*_*i*,tot_ = *Y*_*i*_ + *C*_3_ is the total concentration of component *i, ξ*_*i*_ is its promoter activity in relative promoter units (RPU), *α*_*i*_ is the synthesis rate per RPU in nM/(min RPU), and *δ*_*i*_ is the degradation rate constant in min^*−*1^, for *i* 1, 2 . At steady state, 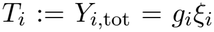, where *g*_*i*_ = *α*_*i*_*/δ*_*i*_. Since we assume that binding equilibrates faster than expression, the component totals are conserved on the binding time scale. So, we set *Ċ*_3_ = 0 and define *K* = *k*_*r*_*/k*_*f*_ to obtain *KC*_3_ = (*T*_1_ − *C*_3_)(*T*_2_ − *C*_3_). Therefore, the equilibrium complex concentration satisfies

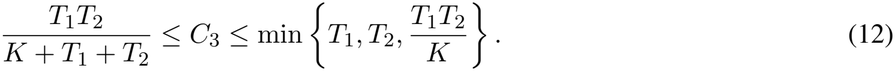

We note that the positive equilibrium satisfies 0 *<C*_3_ < min {*T*_1_, *T*_2_} and we can expand the binding balance to get the lower bound *T*_1_*T*_2_ = (*K* + *T*_1_ + *T*_2_ − *C*_3_)*C*_3_ ≤ (*K* + *T*_1_ + *T*_2_)*C*_3_. The upper bound can be obtained from *KC*_3_ ≤ *T*_1_*T*_2_, and the conservation laws *C*_3_ ≤ *T*_*i*_.

We next establish uniqueness and monotonicity of the equilibrium to justify the output bounds in (10). The balance has a unique solution in (0, min {*T*_1_, *T*_2_} ) because (*T*_1_−*C*_3_)(*T*_2_ − *C*_3_) − *KC*_3_ is positive at zero, negative at the other endpoint, and strictly decreasing on this interval. So, implicit differentiation gives

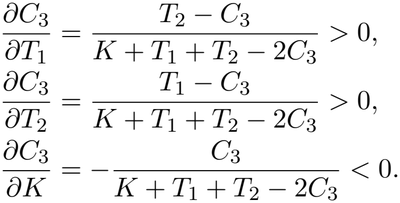

The denominator is positive since *T*_*i*_ *> C*_3_ and *K >* 0. At the selected values of *K* and *k*_out_, the minimum and maximum reporter flux occur at the lower and upper component total endpoints, respectively. These endpoints give the bounds in (10) over the given input intervals.

### B. HrpV inhibition and weaker split T7 association

We use the following reporter relations to bound the output of the HrpV inhibition and weaker split T7 association designs. HrpV binds the HrpR/HrpS activating complex and suppresses reporter production. Under the fast and strong binding assumption, the inhibitor total *V*_tot_ gives the reporter relation

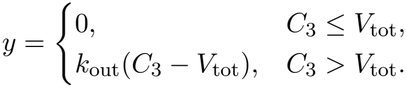

This relation is nondecreasing in *C*_3_. The component total endpoints therefore determine the output bounds for the selected inhibitor total. Weaker split T7 fragment association changes *K* in the binding balance. The same derivatives give the output bounds at the selected value of *K*.

### C. InvF/SicA with SipC

Let *Z* be the SipC concentration and *C*_2*Z*_ the concentration of the SicA and SipC complex. The input production fluxes are *p*_*i*_ = *γ*_*i*_*T*_*i*_, and *p*_*Z*_ is the fixed SipC production flux. The CRN dynamics of this circuit are

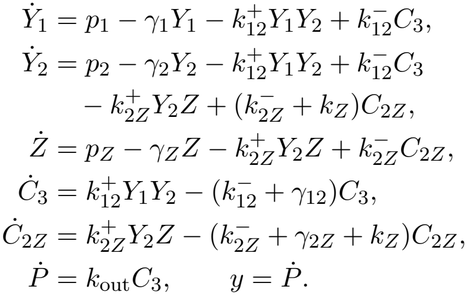

All *γ* parameters are degradation rates, *k*^+^ and *k*^*−*^ are association and dissociation rates, and *k*_*Z*_ is the SipC secretion rate constant [31]. The secretion reaction *C*_2*Z*_ → *Y*_2_ releases SicA. All rates are positive. An equilibrium solution of the concentration balances exists and is unique. This can be seen directly from the equilibrium balance

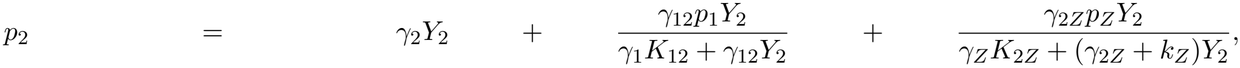

where 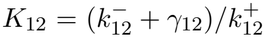 and 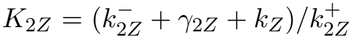. The right hand side is zero at *Y*_2_ = 0, strictly increasing in *Y*_2_, and unbounded as *Y*_2_→ ∞. Therefore it equals *p*_2_ *>* 0 at exactly one positive value of *Y*_2_, which uniquely determines the remaining concentrations and reporter flux. At the equilibrium, the remaining concentrations are

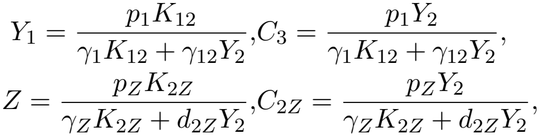

where *d*_2*Z*_ = *γ*_2*Z*_ + *k*_*Z*_. The scalar balance also establishes monotonicity in the input production fluxes. Write the third term on the right hand side as *B*(*Y*_2_) = *γ*_2*Z*_*C*_2*Z*_(*Y*_2_), which has *B*^*′*^(*Y*_2_) *>* 0. For the expression *C*_3_(*p*_1_, *Y*_2_) above, both partial derivatives *∂C*_3_*/∂p*_1_ and *∂C*_3_*/∂Y*_2_ are positive. Then, implicit differentiation gives

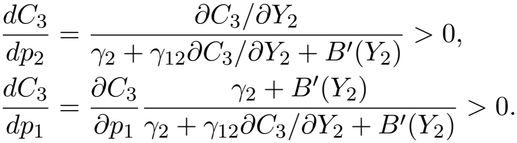

Thus *y* = *k*_out_*C*_3_ increases with each input production flux when the other parameters are fixed. Since *p*_*i*_ = *γ*_*i*_*T*_*i*_, the lower and upper component total endpoints give the corresponding flux bounds in (10).

## Appendix D

Promoter Dynamics for the NAND Circuit

We derive the equilibrium response of each promoter module to establish the component interfaces used in Section VII-C. Regulator binding increases transcription above the basal rate in each activated expression module. Let *D*_0_, *D*_1_ denote vacant and occupied promoter abundance, *A* the regulatory activity, *m* RNA abundance, and *p* the protein output. Binding conserves the total promoter abundance *T* . The association and dissociation constants are *k*_*b*_, *k*_*u*_, the transcription and translation constants are *k*_*t*_, *k*_*ℓ*_, and the RNA and protein degradation constants are *d*_*m*_, *d*_*p*_. These rate constants are positive, and *α*≥ 0 is the basal transcription coefficient. With net binding flux *J*, the model is

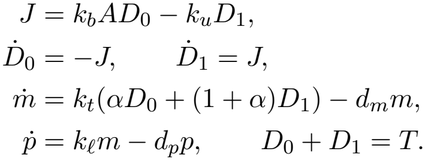

At equilibrium, the binding balance fixes promoter occupancy. With *q* = *k*_*b*_*A/k*_*u*_, the RNA and protein balances give

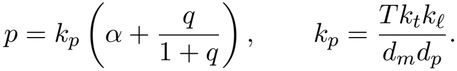

The conservation of promoter gives the saturation denominator, and the production and degradation rates determine the amplitude *k*_*p*_. We model the AND promoter with states *D*_00_, *D*_10_, *D*_01_, *D*_11_, whose indices record occupancy by the two regulatory inputs, with regulatory activities *A*_*R*_, *A*_*S*_. Each input has association and dissociation constants that are independent of occupancy by the other input. The net binding fluxes *J*_*R*0_, *J*_*R*1_ describe binding of the first input when the other site is vacant or occupied, respectively. The fluxes *J*_*S*0_, *J*_*S*1_ describe the corresponding transitions for the second input. The fully occupied state drives production of CI RNA, denoted by *m*_*C*_. We use the same notation for constants with subscript *C*, as before. So, the model is

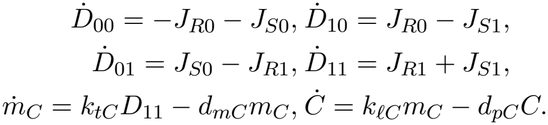

For example, *J*_*R*0_ = *k*_*bR*_*A*_*R*_*D*_00_ − *k*_*uR*_*D*_10_ and *J*_*R*1_ = *k*_*bR*_*A*_*R*_*D*_01_ − *k*_*uR*_*D*_11_. The fluxes for the second input follow the corresponding transitions. For total AND promoter abundance *T*_*A*_, conservation gives *D*_00_ + *D*_10_ + *D*_01_ + *D*_11_ = *T*_*A*_. With *q*_*R*_ = *k*_*bR*_*A*_*R*_*/k*_*uR*_ and *q*_*S*_ = *k*_*bS*_*A*_*S*_*/k*_*uS*_, the equilibrium active promoter fraction and CI level satisfy

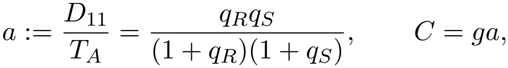

where *a* is the active promoter fraction, and *g* = *T*_*A*_*k*_*tC*_*k*_*ℓC*_*/*(*d*_*mC*_*d*_*pC*_ ) is the CI expression gain.

The CI then represses transcription in the reporter module. To describe this repression, we let *N*_0_, *N*_1_ denote vacant and occupied promoter abundance, with conserved total *N*_0_ + *N*_1_ = *T*_*N*_ . The CI regulatory activity is *A*_*C*_, and the net binding flux is 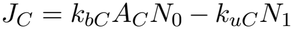. The promoter balances are 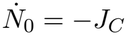 and 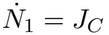. Then, the reporter RNA abundance *m*_*G*_ and fluorescence *G* satisfy the following balances, with rates indexed by *G* and basal transcription coefficient *α*_*G*_:

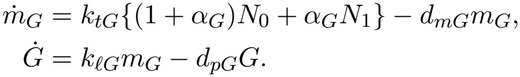

At equilibrium, we define *q*_*C*_ = *k*_*bC*_ *A*_*C*_*/k*_*uC*_ and obtain the reporter response

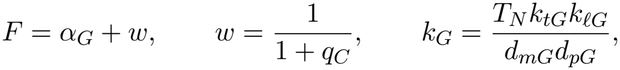

where *w* is the unoccupied reporter promoter fraction. So, the normalized fluorescence decreases from 1 + *α*_*G*_ to *α*_*G*_ as CI regulatory activity increases. For fixed regulatory activities, the binding balances and conservation laws uniquely determine promoter occupancy. The production and degradation balances then uniquely determine the

RNA and protein expression levels. Thus, these models establish unique equilibrium responses and the interface relations used in Section VII-C.

## Acknowledgment

The author thanks the CDC reviewers for helpful suggestions on the conference version of this work and Shruthi Rao, undergraduate researcher at UC Merced, for extracting the Cello library data. The author used OpenAI Codex GPT 6.1 for extracting experimental data from the relevant papers, assistance with computational implementations, and to generate visualizations. All AI outputs were carefully reviewed, vetted, and edited by the author for accuracy and presentation. The author takes full responsibility for the content.

